# A Novel Genome Optimization Tool for Chromosome-Level Assembly across Diverse Sequencing Techniques

**DOI:** 10.1101/2023.07.20.549842

**Authors:** Wei-Hsuan Chuang, Hsueh-Chien Cheng, Yu-Jung Chang, Pao-Yin Fu, Yi-Chen Huang, Ping-Heng Hsieh, Shu-Hwa Chen, Pui-Yan Kwok, Chung-Yen Lin, Jan-Ming Ho

## Abstract

This paper introduces a novel genome assembly optimization tool named LOCLA, which stands for “Local Optimization for Chromosome-Level Assembly”. It identifies reads and contigs aligned locally with high quality on gap flanks or scaffold boundaries of draft assemblies for gap filling and scaffold connection. LOCLA applies to both de novo and reference-based assemblies. It can also utilize reads produced by diverse sequencing techniques, e.g., 10x Genomics (10xG) Linked-Reads, and PacBio HiFi reads.

We validated LOCLA on three human samples and one non-model organism. For the first two human samples, LLD0021C and CHM13, we generated de novo draft assemblies from 10xG Linked-Reads. On LLD0021C, LOCLA improves the draft assembly by adding 23.3 million bases using only 10xG Linked-Reads. These additional bases cover 28,746 protein-coding regions, particularly in pericentromeric and telomeric regions. On the CHM13 sample, we took 10xG Linked-Reads and PacBio HiFi reads as input. As a result, LOCLA added 46.2 million bases to the draft assembly. The increased content enables us to identify genes linked to complex diseases (e.g., ARHGAP11A) and critical biological pathways. We created two reference-guided draft assemblies on the third human sample, HG002, using contigs assembled from PacBio HiFi reads. LOCLA enhances the two draft assemblies by recovering 27.9 million bases (22.26%) and 35.7 million bases (30.93%) of the sequences discarded by the reference-guided assembly tool. The results indicate the robustness of LOCLA’s contig detection algorithm on gap flanks. Furthermore, we show that 95% of the sequences filled in by LOCLA have over 80% accuracy compared with the HG002 reference genome published by the Human Pan-genome Reference Consortium. On the non-model organism, LOCLA enhanced the genome assembly of Bruguiera sexangula (JAHLGP000000000) by decreasing 41.4% of its gaps and raising the Benchmarking Universal Single-Copy Orthologs (BUSCO) analysis score to 98.10%.

LOCLA can optimize de novo and reference-guided assemblies using varied sequencing reads. The final assemblies produced by LOCLA have improved in both quantity and quality. The increased gene content may provide a valuable resource in advancing personalized medicine.

## Introduction

Since the advent of genome-sequencing technology, resolving the assembly of the human genome has been repeatedly attempted for over 20 years. In 2003, the publication of the first near-complete human reference genome marked a triumph in biomedical research^1^. Advances in sequencing techniques, from the initial Sanger Sequencing to next-generation sequencing (NGS), have enabled the high-throughput generation of sequencing data; TGS platforms characterized by long read sequences have paved the way to accomplishing accurate human genome sequencing^2^. The first version of the representative human genome, GRCh38, was released in 2013. The GRCh38 reference genome has hitherto been the basis of human genomic studies for identifying functional genetic regions, defining regulatory elements, comparing genomic diversity, conducting population genomics analyses, and searching for disease-causing mutations^3^. Moreover, the cost of sequencing has decreased, and a substantial amount of human genomes have thus already been sequenced. These sequences have been compared against the human reference genome to identify genetic factors associated with health and disease^4^. The existing practice in genomic analysis is to align the reads generated by sequencers with the GRCh38 reference genome to determine the location of the reads and thus construct the individual’s genome.

An alternative to alignment-based assembly is *de novo* assembly. By rejecting the bias from the reference genome, *de novo* assembly can benefit the genotyping process in two ways: 1) new sequence assemblies for previously unreported genomic regions can be gained, and 2) an individual’s genome can be characterized in a reference-unbiased fashion ^5, 6^. Consequently, gaps in the individual’s genome, relative to the reference, can be resolved and individual variants can be identified rather than being discarded. Increasing the content of personal genomes would enable the identification of unforeseen variants and facilitate disease association tests.

Technologies that provide long-range genomic information with acceptable accuracy should be incorporated into the characterization of large structural mutations in disease diagnosis. Although barcodes are employed to provide long-range information in some synthetic long-read techniques, such as 10x Genomic sequencing (10xG), these techniques inherit the similar limitations as NGS for highly repetitive genomic regions^7^. One possible solution is to incorporate optical mapping (OM) into the sequencing pipeline. If integrated with other sequencing reads, OM indicates the order and orientation of sequence fragments, identifies and corrects potential chimeric joins, and estimates the size of the gap between adjacent reads^8^.

The 10xG Linked-Reads technique uses molecular barcodes to tag reads generated from high-molecular-weight DNA^9^. The key concept is to introduce a unique barcode to every short read derived from a few individual molecules. By tracing the same barcodes, short reads in different fragments can be linked. A genome assembler designed with 10xG, the Supernova assembler, uses read pairs to cover short gaps. Barcodes are also informative for filling in large gaps. If the physical locations of two scaffolds are actually close, multiple molecules in the partitions would be highly likely to bridge the gap between two scaffolds. Linked-Reads can provide long-range information at a length of 100 kbp, which is a major improvement compared with Illumina short-read sequencing with the range information at a length of 300 bp.

In Bionano OM, specific 6-mer sequence motifs are set as markers to provide a blueprint of the genome structure ^8^. The Bionano optical map may be used to order and orient sequence fragments, identify potential chimeric joins, and help estimate the size of the gap between adjacent sequences. To further provide long-range information for disentangling complex genomes, Mostovoy and colleagues^10^ proposed a hybrid method combining Illumina sequencing, 10xG, and Bionano OM to resolve end-to-end, chromosome-level human genome assembly. However, according to their published results, numerous N-base gaps were interspersed in the scaffolds; thus, numerous contigs remained. The hybrid method was applied to assemble 17 human genomes from five populations in another study, and the comprehensiveness of the assembled genome resulted in the discovery of thousands of non-reference unique insertions^11^. These results challenged the representative human genome, indicating that it did not include some common genomic structures. By considering more than 300 human samples collected from diverse populations, Wong and colleagues^4^ constructed a human diversity reference genome (HDR) by using the same hybrid pipeline with the addition of PacBio assemblies, which incorporates non reference unique insertions into the linear reference genome structure. The HDR has considerably improved annotations and interpretation of structural variants that were not previously approachable due to fragmented scaffolds. It has also increased the accuracy of the alignment-based variant caller while retaining high efficiency due to the linear genome structure. On the basis of the reviewed studies, we are convinced that the complementarity of Linked-Reads and optical maps has high potential to make the production of higher quality genomes more routine and economical.

In our study, we designed a genome assembly optimization tool that depends primarily on long-range genomic information to iteratively fill gaps and extend scaffold lengths. We hereby introduce LOCLA, short-hand notation for “Local Optimization for Chromosome-Level Assembly” The tool suite comprises four main modules: (1) local-contig-based (LCB) gap filling, (2) global-contig-based (GCB) gap filling, (3) GCB scaffolding, and (4) LCB scaffolding. The basic concept of each method is presented in Figure 1. In (1), contigs generated from short reads with long-range information, i.e., contigs assembled from 10xG Linked-Reads, which are named “local-contigs” here, are used to fill the gaps in a scaffold. In (2), TGS long reads or scaffolds, which are named “global-contigs” here, produced using other sequence assemblers are used to fill gaps in a scaffold. In (3) and (4), the reads are used to further extend the existing scaffolds to span a much wider range.

**Figure 1.**
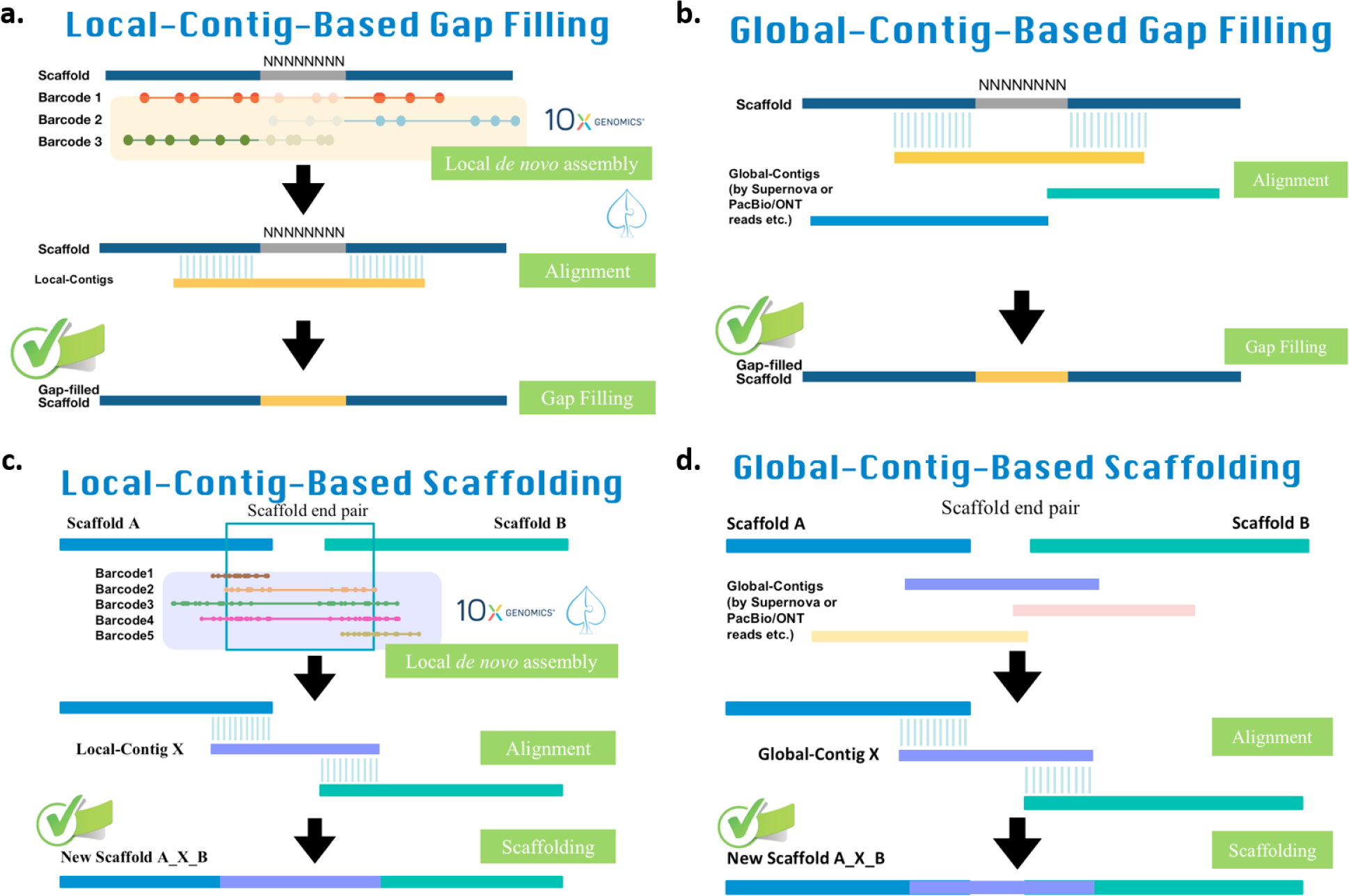
| Core concept of the four main LOCLA modules. **a,** “Local-Contig-Based (LCB) Gap Filling”: First, we align all barcoded linked-reads to the scaffolds and de novo assemble Local-contigs using the reads belonging to barcodes mapped within gap flanks. Then, we map these contigs onto the scaffolds and determine the best hit to fill in gaps. **b,** “Global-Contig-Based (GCB) Gap Filling”: We align Global-contigs to all scaffolds and find the best hit to fill in gaps. **c,** “Local-Contig-Based (LCB) Scaffolding”: We align all barcoded linked-reads to the head and tail of scaffolds and pair up scaffolds with shared barcodes. Then we construct Local-contigs from the barcoded-reads and connect scaffolds with the optimal L-contig. **d,** “Global-Contig-Based (GCB) Scaffolding”: Identical to LCB Scaffolding, we align linked-reads to the ends of scaffolds and pair up scaffolds with shared barcodes. Global-contigs are then mapped onto all scaffold pairs and connected with the most ideal G-contig.

The 10xG official assembler Supernova was used as a benchmark to demonstrate the efficacy of LOCLA on three human samples, LLD0021C [Data Citation 1] and CHM13^12^. For LLD0021C, we adopted the aforementioned hybrid method of 10xG Linked-Reads and Bionano Genomics OM and discovered that the N50 value of the Supernova assembly increased from 45 to 59 Mbp. LOCLA added an additional 23 Mbp and closed 9,700 gaps in the Supernova assembly. With these additional bases, we identified 136 functional genes related to complex diseases and biological pathways that were obscured in the Supernova assembly. For CHM13, LOCLA successfully increased N50 from 39 to 44 Mbp and increased the total genome size by 38 Mbp. We improved the resolution of 145 functional genes associated with pathological findings on the Supernova assembly and discovered 10 exclusive genes. We also observed that LOCLA performed well in the assembly of regions that are considered difficult to solve or involve repetitive sequences (i.e., higher resolution was achieved for 3,768 noncoding transcripts and 2,996 pseudogenes). For HG002, LOCLA successfully filled in a large number of gaps in the RagTag assemblies, with 2,877,149 and 2,346,125 gaps filled respectively. Furthermore, LOCLA was able to retrieve a significant number of contigs that RagTag did not use, with 22.26% for the GUA assembly and 30.93% for the GMA assembly. This highlights the effectiveness of LOCLA’s candidate contig detection algorithm on gap flanks. The GMA assembly contains more information than the GUA assembly due to its retention of multiple-aligned sequences, which allowed LOCLA to locate and recruit more candidate contigs for further gap filling.

LOCLA has also demonstrated its effectiveness in improving the quality of genome assembly by utilizing 10xG Linked-Reads in another example, the *Bruguiera sexangula* dataset. Pootakham W. *et al.*^69^ performed the assembly of the Bruguiera sexangula genome, which contains 260,518,658 base pairs and 1,627,214 gaps. LOCLA filled in 674,896 gaps, which is equivalent to 41.4% of the total gaps in the initial assembly. In addition, LOCLA enhanced the draft assembly by incorporating an additional 7,404,783 bases using 10xG Linked-Reads. As a result, the BUSCO score increased from 97.90% to 98.10%.

## Results

In the following text, we demonstrate how LOCLA was used to improve the assembly quality of four genome samples. The first three are human samples, i.e., LLD0021C, CHM13 and HG002 data sets, respectively. The fourth is a non-model organism, *Bruguiera sexangula*.

LOCLA is an optimization tool that improves the quality of draft assembly by iteratively filling gaps and extending scaffolds. From the results, we prove that an increase in gene content leads to a clearer view of genetic information and enables further insight into functional genes, especially those related to diseases. We also demonstrated that LOCLA is flexible using either 10xG Linked-Reads or TGS sequencing reads.

### LOCLA assembly of LLD0021C compared with Supernova assembly

The LLD0021C data set comprises the results of 10xG Linked-Reads and Bionano OM. A draft was generated by Supernova and was then input to the Bionano Hybrid Scaffold pipeline. The LOCLA submodules were run in order, i.e., GCB gap filling, LCB gap filling, LCB scaffolding and finally GCB scaffolding, to produce the final assembly. For LLD0021C (Table 1), LOCLA filled in 23,319,370 new base pairs in the 9777 gaps that were present in the Supernova draft (Supplementary Table 5). Among these gaps, 5785 were completely filled, and 3992 were partially filled. In our experiment, the mean length of the completely filled gaps was 113.73bp, and the mean length of the partially filled gaps was 4,274.27 bp. In addition, the largest gap size for each type of gap was 59,608 and 100 kbp, respectively, indicating that LOCLA is capable of mending large gaps. We also raised N50 from 45,208,438 bp to 59,229,662 bp, an increase of 14 Mbp. The maximum scaffold length was increased to approximately 130 Mbp, longer than 12 pairs of human chromosomes and approximately the length of chromosome 11 (135,186,938 bp). To demonstrate that LOCLA can achieve higher resolution than previous methods in functionally important genomic regions, we extracted sequences that were filled exclusively by LOCLA. The LOCLA-filled sequences were then annotated on the basis of GENCODE in GRCh38 coordinates^16^. We classified the LOCLA-filled sequences into four main GENCODE biotypes: coding sequences, noncoding transcripts, pseudogenes, and others (Table 2). We discovered that LOCLA retained sequences in more than 28,000 protein coding regions, indicating that these sequences with direct functional impact were missing in the Supernova assembly. Specifically, LOCLA significantly improved the genome content of genes located in pericentromeric and telomeric regions, including sequences in exons and transcripts (Table 3).

**Table 1.**
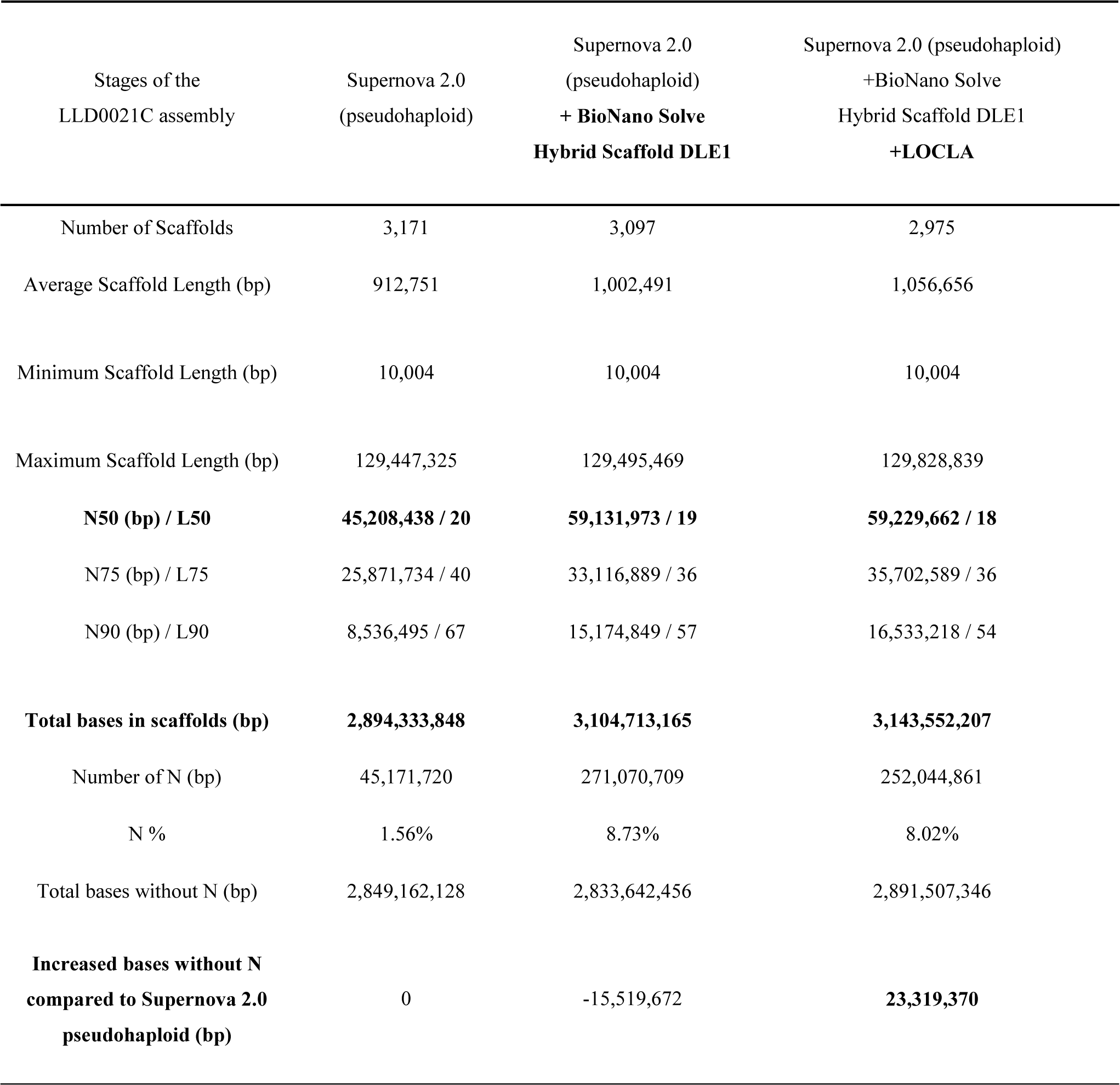
| Assembly statistics show notable increase especially in N50 and total base length on sample LLD0021C after LOCLA.

**Table 2.** | LOCLA-filled content in the Supernova assembly categorized by GENCODE biotype (sample:LLD0021C)

| **Biotype** | **Counts of Filled Genomic Regions** |
| --- | --- |
| Coding Sequence: | 28,861 |
| Protein coding | 28,746 |
| Immunoglobulin and T cell receptor | 115 |
| Non-coding Transcript: | 3,768 |
| Long non-coding RNA (lncRNA) | 3,552 |
| Non-coding RNA (ncRNA) | 206 |
| tRNA | 7 |
| rRNA | 3 |
| Pseudogene: | 2,996 |
| Unprocessed pseudogene | 2,328 |
| Processed pseudogene | 504 |
| Unitary pseudogene | 78 |
| rRNA pseudogene | 9 |
| Other pseudogene | 77 |
| Others | 44 |
| To be experimentally confirmed (TEC) | 44 |

**Table 3.** | LOCLA significantly improves genome contents in protein coding genes located in pericentromeric, telomeric regions, and difficult-to-solve regions. (sample:LLD0021C)

| Gene name | Filled gene length (bp) | Filled transcript count | Filled exon count | Genomic Position |
| --- | --- | --- | --- | --- |
| TPTE2P6 | 76,390 | 9 | 3 | Chr13 pericentromeric |
| PARP4 | 63,621 | 12 | 34 | Chr13 pericentromeric |
| MAD1L1 | 62,662 | 45 | 13 | Chr7 telomeric |
| PLD5 | 58,127 | 37 | 10 | Chr1 telomeric |
| PKD1 | 38,435 | 32 | 46 | Chr16 telomeric |
| PDPK1 | 38,232 | 26 | 14 | Chr16 telomeric |
| TRIM16 | 36,321 | 19 | 12 | Chr17 pericentromeric |
| DDX11 | 30,947 | 60 | 27 | Chr12 pericentromeric |
| RASA4B | 35,309 | 10 | 21 | Chr7 Overlap POLR2J3 |
| POLR2J3 | 25,522 | 24 | 7 | Chr7 Overlap RASA4B |

For noncoding transcription, 3552 additional sequences in lncRNA were identified in the LOCLA assembly. lncRNAs have long been regarded as key regulatory elements for gene expression. LOCLA could resolve three sequences encoding rRNA, which are the most difficult-to-solve regions in genome assembly problems. LOCLA also improved the genome content of approximately 3000 pseudogenes. Pseudogenes are sequences generated through genome duplication and retrotransposition in the evolutionary process. Therefore, pseudogenes comprise duplicated and repeated sequences that are considered difficult to assemble. In general, LOCLA outperformed Supernova in multiple functional classes.

### Evaluating LOCLA assembly of LLD0021C with respect to the standard human reference, GRCh38

For evaluation, we aligned the Supernova assembly of LLD0021C before and after performing LOCLA to the latest representative human genome, GRCh38.p13, using *minimap2*^20^. We kept the mapped sequence alignments first, then applied two filter criteria on the alignments: mapping quality (MQ) equal to 60 and mapping identity (MI) over 70%. A score of 60 in MQ represents the accuracy of the alignment position, while a score of 70% in MI exhibits the high resemblance between sequences, for it is calculated by dividing the length of sequence matches by the sum of the lengths of the query and deletions. From the results shown in Table 5, we see that the number of scaffolds and percentage applying each filter criterion increased after performing LOCLA. At the same time we proved that among the 23 Mbp LOCLA added to the Supernova assembly, around 20 Mbp has high quality, i.e., MQ=60 and MI >=70%.

Only 8 out of the 2975 scaffolds in the LOCLA assembly were not mapped onto the reference; 155 out of 3171 scaffolds were not mapped in the Supernova assembly. Because LLD0021C is the genome sample of a Taiwanese person whereas GRCh38 originates from 11 other individuals (approximately 70% of GRCh38 is from just one man), we speculate that the lack of diversity in the reference may be the reason for these unmapped sequences. We believe that these unmapped scaffolds might lead to new findings. Thus, we employed AUGUSTUS^17^ for gene prediction and subsequently used protein BLAST^18^ to investigate whether the predicted sequences are conserved in organisms. Consequently, we identified 11 inferred genes in the sequences. Among these genes, two genes were homologs of the existing genes *PTZ00395* and *DUX4*. Specifically, *DUX4* is located within a repeat array in the sub-telomeric region of chromosome 4q; a similar repeat array is present on chromosome 10.

### LOCLA outshines Supernova in masked regions (N in reference genome) and repeat regions of the GRCh38 reference

Genomic regions marked with a gap of “N” still exist in the GRCh38 human reference genome, especially in homologous centromeres and genomic repeat arrays. These regions with repeated sequence are notorious for their poor resolution in alignment-based short-read *de novo* assembly techniques. For the human sample LLD0021C, LOCLA was able to extend the scaffolds to fill genomic contents in gap regions, accounting for 462,705 bp in 12,046 gap regions, and was discovered to perform significantly better than Supernova did. Assembling contigs and scaffolds in highly repeating regions has been prone to error in *de novo* genome assembly. However, repeat elements comprise a considerable percentage (approximately 45%) of the *Homo sapiens* genome. Therefore, we assessed the performance of LOCLA with the human sample LLD0021C in these repeat regions. We used RepeatMasker ^19^, which is based on Repbase, to identify repeat regions. Compared with the Supernova assembly, the LOCLA assembly contained 11,031,487 additional bases masked and identified as repeat elements; 1,138,402 and 5626 bp were classified into the short and long interspersed nuclear element categories, respectively (Supplementary Table 6). Surprisingly, 218 repeat patterns were identified in the LOCLA assembly but not in Supernova.

### LOCLA on the CHM13 cell line

Unlike LLD0021C, the CHM13 data set comprises only 10xG Linked-Reads; the Bionano Hybrid Scaffold pipeline ^8^ is not included. The LOCLA assembly contained 46,287,195 more base pairs than were present in the initial Supernova haplotype and reduced the number of gaps from 35,124,040 to 25,648,758 (Table 4). Moreover, the maximal scaffold length and N50 had increased by 11.5 and 5 Mbp, respectively. Upon further examination, we filled in 18,636 of the 23,349 gaps; 10,768 were completely filled and 7868 were partially filled (Supplementary Table 7). Similar to the results for LLD0021C, the lengths of the completely filled gaps were mostly <1 kbp; however, the largest completely filled gap was of approximately 100 kbp. In summary, even without Bionano OM, LOCLA could still produce useful results and enhance the assembly quality.

**Table 4.** | LOCLA effectively expands total genome size and reduces gaps on CHM13 assembly.

| Stages of the CHM13 assembly | Supernova 2.1 (pseudohaploid) | Supernova 2.1 (pseudohaploid) + LOCLA |
| --- | --- | --- |
| Number of Scaffolds | 4,999 | 4,809 |
| Average Scaffold Length (bp) | 583,041 | 613,731 |
| Minimum Scaffold Length (bp) | 10,000 | 10,000 |
| Maximum Scaffold Length (bp) | 91,225,906 | 102,752,858 |
| **N50 (bp) / L50** | **39,045,223 / 25** | **44,037,625 / 23** |
| N75 (bp) / L75 | 18,387,696 / 51 | 19,878,974 / 48 |
| N90 (bp) / L90 | 1,578,658 / 108 | 1,799,962 / 101 |
| Total bases in scaffolds (bp) | 2,914,622,463 | 2,951,434,376 |
| Number of N (bp) | 35,124,040 | 25,648,758 |
| N % | 1.21% | 0.87% |
| Total bases without N (bp) | 2,879,498,423 | 2,925,785,618 |
| **Increased bases without N compared to Supernova 2.1 pseudohaploid (bp)** | **0** | **46,287,195** |

**Table 5.**
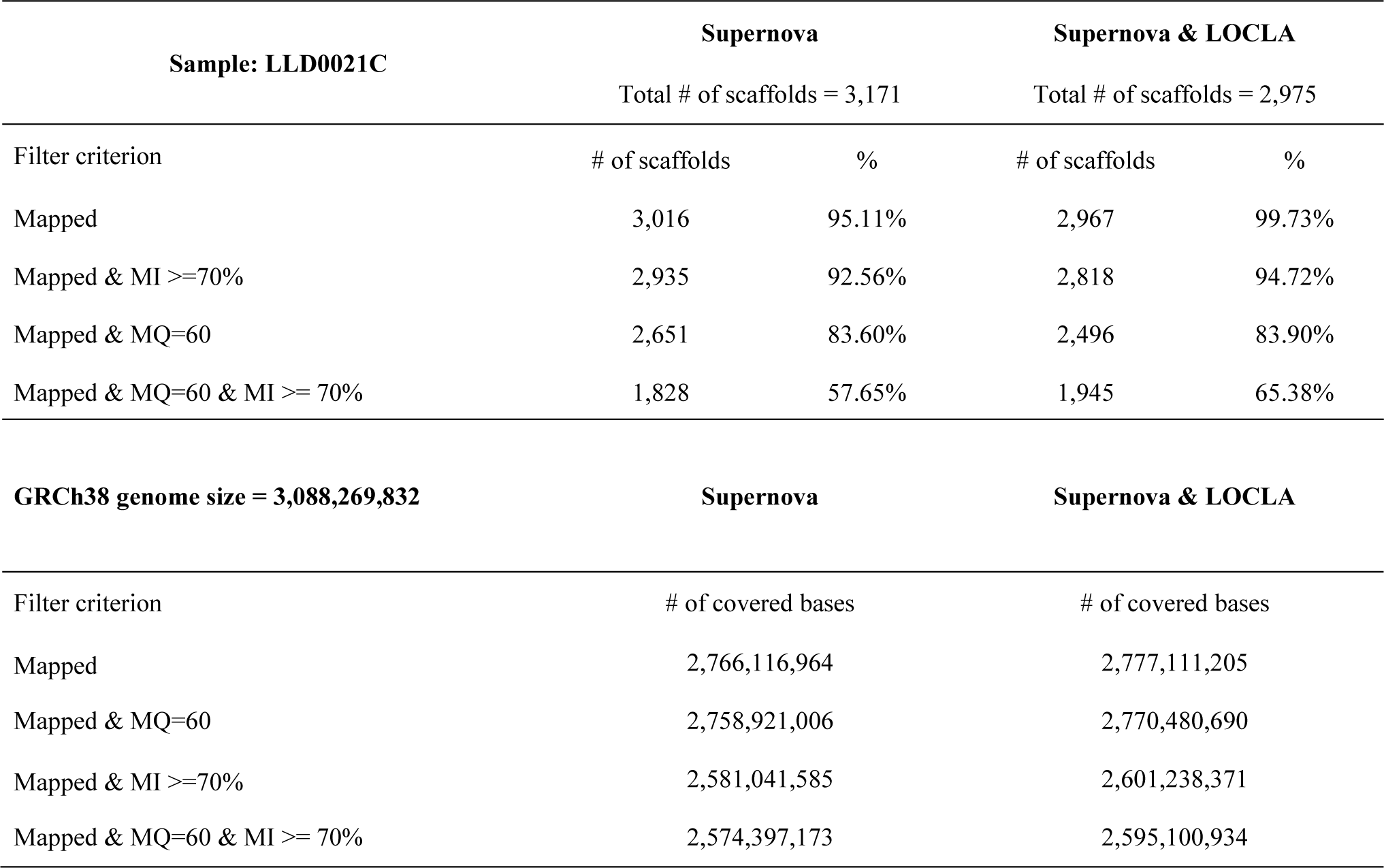
| Evaluation of LLD0021C on the reference genome GRCh38.

### Evaluating LOCLA assembly of CHM13 on the complete human reference genome

To perform an evaluation, we aligned both the Supernova and LOCLA assemblies with the reference genome CHM13v1.1^12^ by using *minimap2*^20^. It is the complete sequence of a human genome constructed by the Telomere-to-Telomere (T2T) Consortium; the genome is available from the National Center for Biotechnology Information (NCBI) as GenBank assembly accession: GCA009914755.3. We applied the same criteria used on LLD0021C to filter alignments. The results presented in Table 6 reveal that the 118 unmapped scaffolds of the Supernova assembly were mapped onto the reference genome after we performed LOCLA, this manifests LOCLA’s capability to correct sequences. It is also evident that the number of scaffolds and covered bases of the assembly all increased with the aid of LOCLA.

**Table 6.**
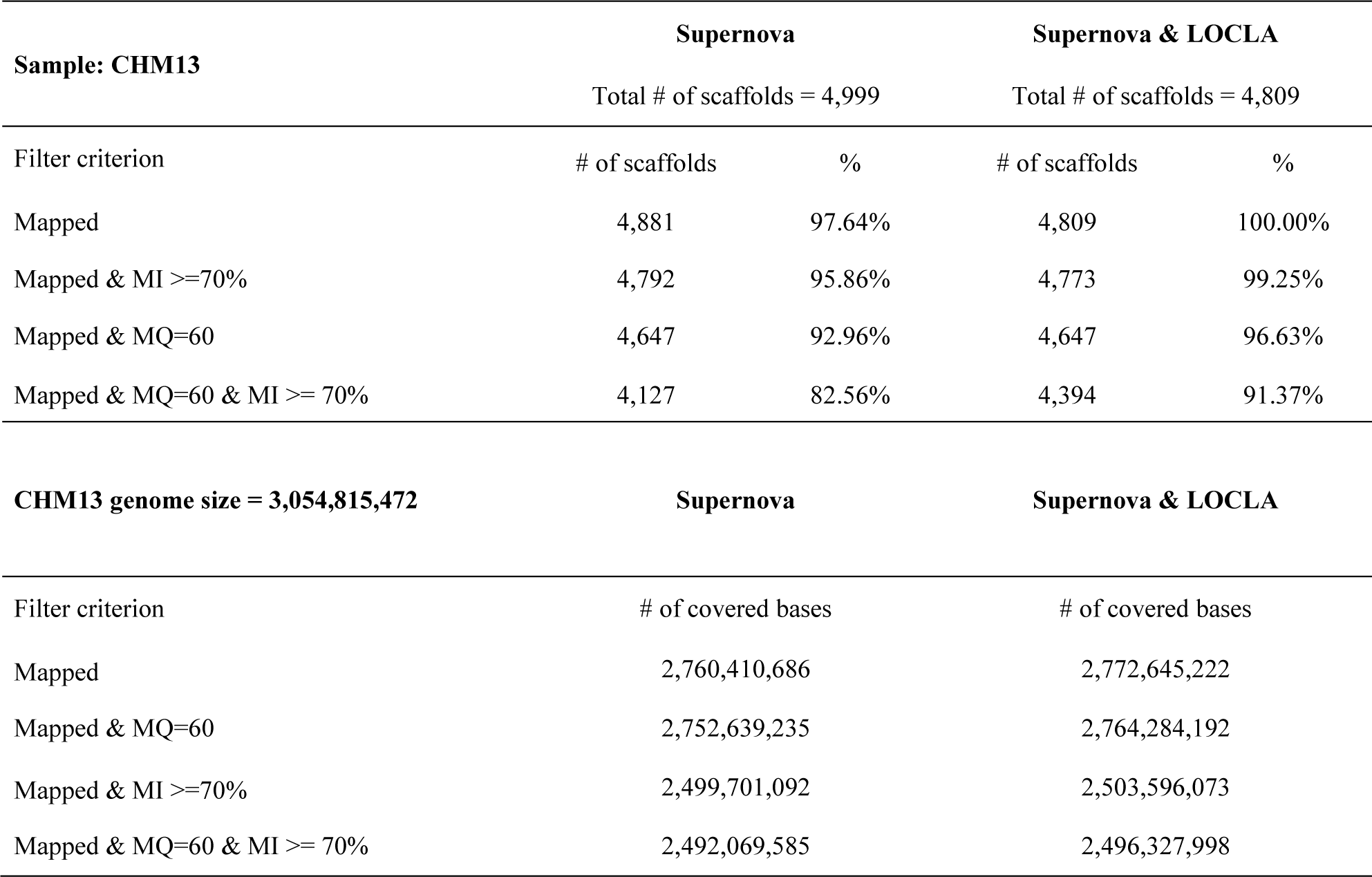
| Evaluation of CHM13 on the reference genome CHM13v1.1.

The LOCLA results were evaluated with the aforementioned pipeline of annotations and identification of repeat elements. Again, LOCLA had higher performance in all functional classes and repeat elements than did Supernova; LOCLA achieved 9.1% higher genomic content than Supernova in CHM13. Notably, LOCLA achieved a 37.73% increase in exon regions in CHM13 and 6.69% increase in ncRNA regions. The CHM13 genome is an effectively haploid genome and has a relatively high proportion of disease-related mutations. We discovered that the LOCLA assembly was a considerable improvement over the Supernova assembly for CHM13.

### LOCLA’s contribution in functional analysis of LLD0021C and CHM13

The LOCLA assembly can significantly improve the quality of functional analysis of an individual human genome and thus provide insights regarding health care. Filling gaps increases gene content related to complex diseases or involved in important biological pathways, including functions such as DNA repair, DNA replication, cell cycle checkpoints, cell signaling transduction, and telomere regulation. These genes are related to cell over-proliferation and tumor development, and they may enable an explicit interpretation of disease mechanisms. LOCLA was discovered to increase the content for each of these genes by hundreds to thousands of base pairs for both LLD0021C and CHM13.

For LLD0021C, LOCLA improved the resolution of 136 genes that were unclear on the Supernova assembly. Regarding *DDX11*, LOCLA filled an additional 31 kbp compared with the Supernova assembly. *DDX11* is involved in DNA replication^21^, DNA repair^22^, heterochromatin organization^23^, and cell cycle regulation and interacts with genes related to meiosis and cell cycle checkpoints^24^, including *STAG1*^25^, *STAG2* ^26^, *SMC1A* ^27^, *CHTF18* ^28^, and *DDX11-AS1*^29^ (Supplementary Table 15). These genes have been reported to be direct disease-causing factors for several cancers, including breast, colorectal, prostate, and gastric cancers. The LOCLA assembly process also enabled us to distinguish the reads of their paralogs. Compared with Supernova, LOCLA increased the gene content in *RTEL1*, one of *DDX11*’s paralogs. *RTEL1* is functionally important in telomere regulation during tumor development and is located in the telomere region of chromosome 16. Chromosome telomeres are difficult to analyze not only by using alignment-based methods but also when using *de novo* genome assemblers ^30, 31^. We inferred that because reads of *DDX11* and its paralogs are largely identical in short genomic ranges, LOCLA may have assembled them accurately. Moreover, in the analysis based on the GRCh38 human reference genome, LOCLA identified more than 0.4 Mbp of novel sequences in gap regions and found an additional 11,031,487 bp of repeat sequences in 1168 repeat patterns identified with the Repbase database ^32^. Specifically, 218 repeat patterns were not identified with Supernova.

For CHM13, LOCLA increased the content of 155 genes; 145 of these were present in the Supernova assembly, but 10 were exclusively present in the LOCLA assembly (Supplementary Table 16). These genes are related to not only complex diseases but also the fundamental mechanisms of the human body. For example, the *ARHGAP11A* gene encodes a member of the Rho GTPase-activating protein family, which causes cell cycle arrest and apoptosis. Studies have demonstrated that the disruption of apoptosis may increase cancer invasiveness during tumor progression, stimulate angiogenesis, deregulate cell proliferation, and interfere with differentiation ^33^. *ARHGAP11A* is highly expressed in colon cancers and a human basal-like breast cancer cell line. The gene is also known to be directly linked to Chromosome 15Q13.3 Deletion Syndrome and Prader–Willi syndrome; an intronic variant of this gene may be associated with sleep duration in children.

### LOCLA outperforms other gap-filling methods

Among the plethora of existing gap-filling methods, we demonstrate that LOCLA can mend larger sequence gaps than some well-known gap-filling tools using paired-end reads. One experiment comparing the gap closure results of GapFiller ^64^ and SOAPdenovo ^34^ (Supplementary Table 8) revealed that the average gap lengths closed by the two methods are 264.87 and 148.39 bp, respectively, on GRCh37 chromosome 14. By comparison, the average gap length closed by GABOLA was found to be 1,812.52 bp on LLD0021C (Supplementary Table 5) and 2162.67 bp on CHM13 (Supplementary Table 7). The percentage of the total gap length closed by LOCLA on CHM13 was 84.37%, which exceeds those of both GapFiller and SOAPdenovo by a large margin.

### LOCLA improves the HG002 assembly based on PacBio HiFi reads

We showed that LOCLA delivers great results even without using 10xG Linked-Reads. We chose the HG002 dataset on account of the abundant data made publicly available by The Genome in a Bottle Consortium (GIAB)^70^. First, we generate a draft assembly via Canu^15^ using the entire PacBio HiFi dataset (255 Gbp with 85.1x coverage) released by the Human Pan-genome Reference Consortium (HPRC)^71^ and we obtained a draft assembly containing 1,602 scaffolds (Supplementary Table 17). Afterwards, we performed RagTag on the Canu assemblies with CHM13 v1.1 as reference first. Due to the fact that the default settings of RagTag discards contigs that have multiple alignments on the reference genome, we adopted two approaches for this step. The first is done by following the default settings (minimum mapping quality threshold = 10) of RagTag, while the second is done by eliminating the minimum mapping quality threshold so that multiple-aligned sequences could be recruited. This process yields two RagTag assemblies which we termed the “Globally-Unique-Aligned (GUA)” assembly and the “Globally-Multiple-Aligned (GMA)” assembly. We then filled gaps via LOCLA on both assemblies with the contigs that weren’t utilized by RagTag for scaffold construction. From Table 7, we see that LOCLA filled in 2,877,149 gaps of the GUA assembly and 2,346,125 gaps of the GMA assembly. During the gap filling process, LOCLA retrieved 22.26% of the contigs unused by RagTag for the GUA assembly, while retrieving even more for the GMA assembly (30.93%) as shown in Table 8. This outcome indicates that the candidate contig detection algorithm on gap flanks is the main advantage of LOCLA. By retaining multiple-aligned sequences, the GMA assembly holds more data compared to the GUA assembly. This enables LOCLA to identify and locate candidate contigs for additional gap filling.

**Table 7.** | Status of the HG002 GUA and GMA assemblies after each stage of process.

| Assembly Name | GUA |  | GMA |  |
| --- | --- | --- | --- | --- |
| Process | RagTag | RagTag & LOCLA | RagTag | RagTag & LOCLA |
| Number of Scaffolds | 23 | 23 | 23 | 23 |
| Minimum Scaffold Length | 39,760,224 | 39,760,339 | 41,606,408 | 42,815,186 |
| Maximum Scaffold Length | 257,977,679 | 259,715,312 | 266,182,842 | 267,323,777 |
| N50 (bp) | 155,468,333 | 157,514,529 | 141,664,663 | 146,222,511 |
| L50 | 8 | 8 | 8 | 8 |
| N75 (bp) | 109,686,031 | 110,063,149 | 114,977,456 | 118,349,064 |
| L75 | 14 | 14 | 14 | 14 |
| N99 (bp) | 39,760,224 | 39,760,339 | 41,606,408 | 42,815,186 |
| L99 | 23 | 23 | 23 | 23 |
| Total size (bp) | 3,112,823,811 | 3,123,350,734 | 3,163,409,713 | 3,186,199,263 |
| Number of N (bp) | 14,780,415 | 11,903,266 | 27,480,021 | 25,133,896 |
| Percentage of N | 0.47% | 0.38% | 0.86% | 0.78% |
| **Total size without N (bp)** | **3,098,043,396** | **3,111,447,468** | **3,135,929,692** | **3,161,065,367** |

**Table 8.**
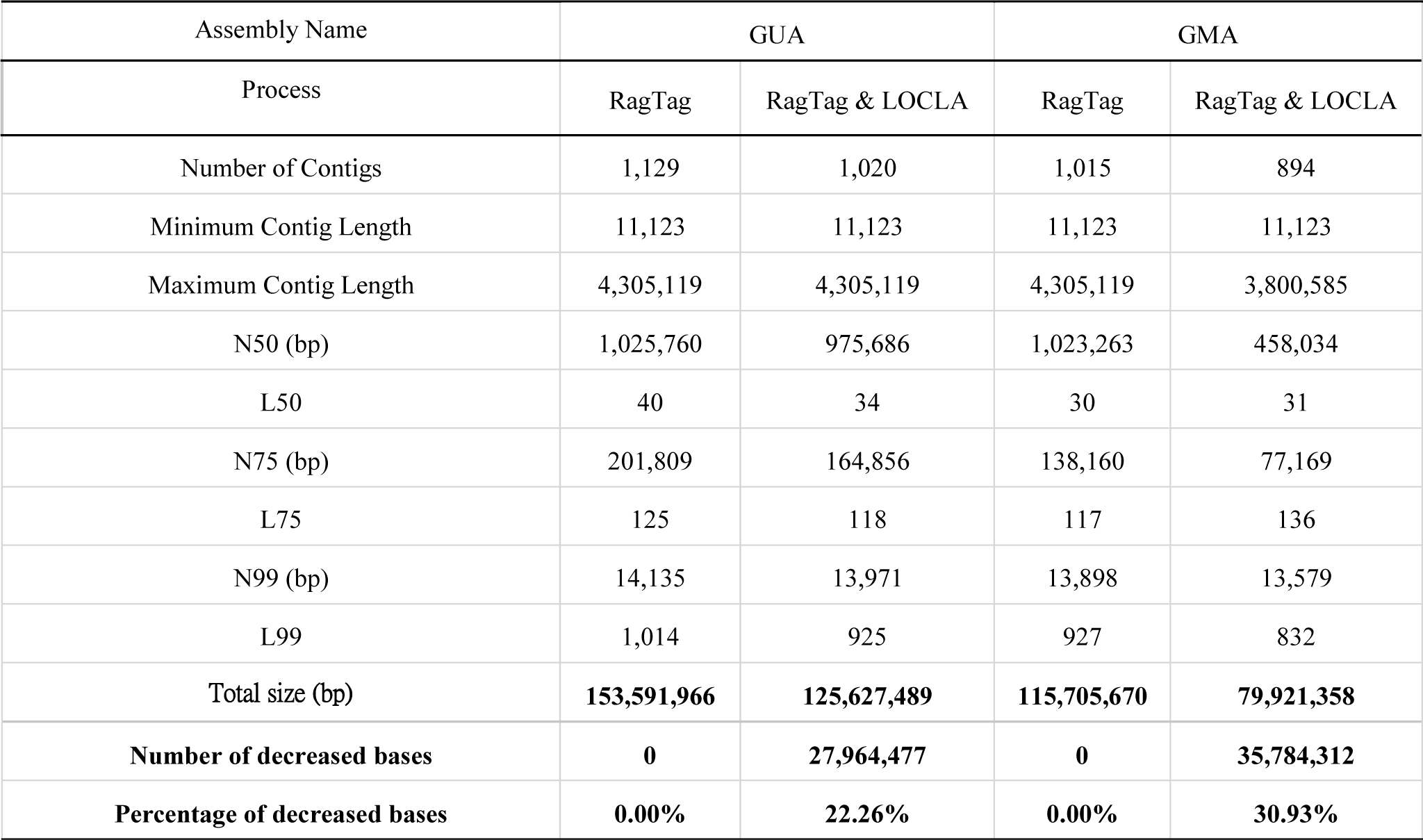
| Remaining contigs after the process of RagTag and LOCLA. (sample:HG002)

| Assembly Name | GUA |  | GMA |  |
| --- | --- | --- | --- | --- |
| Process | RagTag | RagTag & LOCLA | RagTag | RagTag & LOCLA |
| Number of Contigs | 1,129 | 1,020 | 1,015 | 894 |
| Minimum Contig Length | 11,123 | 11,123 | 11,123 | 11,123 |
| Maximum Contig Length | 4,305,119 | 4,305,119 | 4,305,119 | 3,800,585 |
| N50 (bp) | 1,025,760 | 975,686 | 1,023,263 | 458,034 |
| L50 | 40 | 34 | 30 | 31 |
| N75 (bp) | 201,809 | 164,856 | 138,160 | 77,169 |
| L75 | 125 | 118 | 117 | 136 |
| N99 (bp) | 14,135 | 13,971 | 13,898 | 13,579 |
| L99 | 1,014 | 925 | 927 | 832 |
| Total size (bp) | **153,591,966** | **125,627,489** | **115,705,670** | **79,921,358** |
| **Number of decreased bases** | **0** | **27,964,477** | **0** | **35,784,312** |
| **Percentage of decreased bases** | **0.00%** | **22.26%** | **0.00%** | **30.93%** |

### Evaluating LOCLA assembly of HG002 on the HPRC reference genome

For validation, we compared the GUA and GMA assemblies before and after undergoing LOCLA with the maternal haploid assembly published by HPRC. As shown in Supplementary Figure 6, all assemblies have over 91 percent of the genome aligned perfectly to the reference genome, while the percentages of GMA assemblies exceeds the GUA assemblies by a margin. Supplementary Figure 7 illustrates that LOCLA increased the percentage of highly-matched alignments (over 75% of the alignment is matched to the reference) from 93.52% in the RagTag GUA assembly to 96.12% on the GMA LOCLA-optimized assembly. To verify that the sequences LOCLA had filled in are accurate, we performed local alignment on all filled-in sequences, which we will term “patch” in the following text. Figure 2a shows that among the 129 patches, 123 of them are highly similar (mapping identity over 80%) to the reference genome. We selected chromosome 1 and chromosome 13 as an example for a closer look. In Figure 2b and 2c, we see that on chromosome 1, a patch with 5901 base pairs is located within a global alignment with a high mapping identity score (91%). On chromosome 13, a patch is located within an alignment with a lower score (22%) and on the p-arm. Figure 2d and 2e are the local alignment results of these two patches using BLAST. We see that the patch on chr1 (Figure 2d) is aligned to its reference sequence with high quality. While the patch on chr13 has repeatedly aligned to the target sequence due to the innate structure of acrocentric chromosomes, we still see a perfect match among these alignments. This outcome indicates that even in genomic regions containing numerous repeats, LOCLA could still fill in high quality sequences. In Figure 2a, we also noticed that 13 patches are longer than 100kbp and are interested in the accuracy of these patches. All 13 patches have a mapping identity over 79% while 6 of them are over 90%. These findings exhibit the stability in the performance of LOCLA without 10xG Linked-Reads.

**Figure 2.**
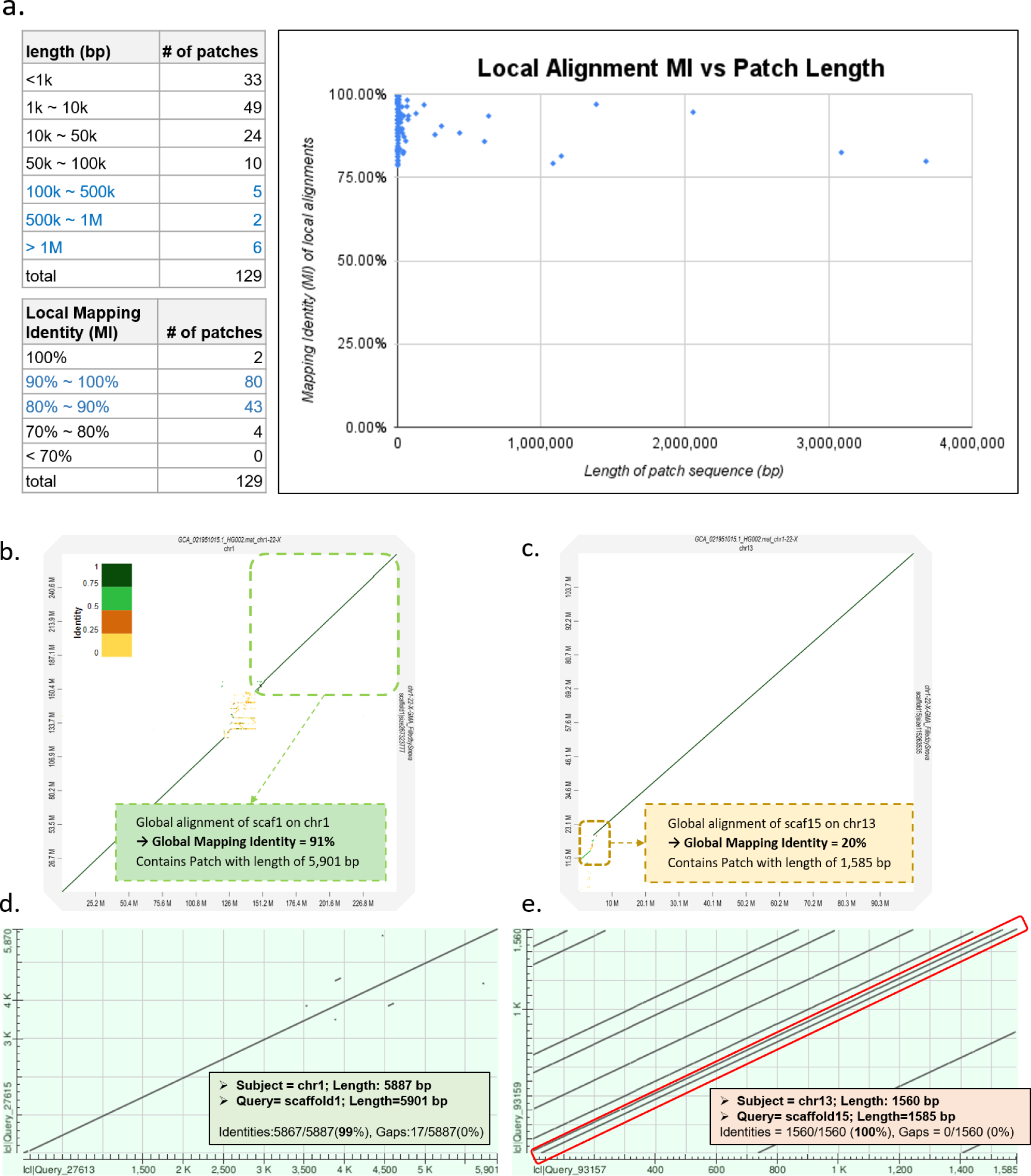
| Evaluating the accuracy of HG002 assembly based on local alignments of patch sequences on the HG002 reference. **a,** The distribution of local alignment Mapping Identity and patch lengths. **b,** The dotplot on shows the alignment between scaffold 1 of the gap-filled GMA assembly and chromosome 1 of the HPRC maternal haploid assembly. Here we focus on a patch located within a global alignment with a high MI score of 91%. **c,** The dotplot on shows the alignment between scaffold 15 of the gap-filled GMA assembly and chromosome 13 of the HPRC maternal haploid assembly. Here we focus on a patch located within a global alignment with a low MI score of 20%. **d,** BLAST results of the first patch with a high global MI (91%) on chr1. We see that the patch is highly similar to its reference sequence. **e,** BLAST results of the second patch with a low global MI (22%) on chr13. Due to the structure of p-arms on acrocentric chromosomes, it is expected that the patch would be repeatedly aligned to the reference. Among the repetitive alignments, there is still a perfectly matched alignment as circled in the plot above.

### Optimization of *Bruguiera sexangula* genome assembly by LOCLA

The *Bruguiera sexangula* genome assembly by Pootakham W. *et al*^69^ has the size of 260,518,658 base pairs containing 1,627,214 gaps. LOCLA filled in 674,896 gaps (41.4% of the number of gaps in the initial assembly) and increased 7,404,783 additional bases to the draft assembly using the 10xG Linked-Reads (Table 9). The BUSCO score was also raised from 97.90% to 98.10%.

**Table 9.** | LOCLA improves the *Bruguiera sexangula* genome assembly through LOCLA.

| Steps | 10xG & RagTag Scaffolding | 10xG & RagTag Scaffolding<br>& LOCLA |
| --- | --- | --- |
| Number of Scaffolds | 20,644 | 20,644 |
| Average Scaffold Length (bp) | 12,620 | 12,946 |
| Minimum Scaffold Length (bp) | 300 | 300 |
| Maximum Scaffold Length (bp) | 17,482,559 | 17,854,050 |
| N50 (bp) / L50 | 11,020,310 / 10 | 11,501,059 / 10 |
| N75 (bp) / L75 | 7,984,485 / 17 | 8,321,598 / 17 |
| Total bases in scaffolds (bp) | 260,518,658 | 267,248,545 |
| Number of N (bp) | 1,627,214 | 952,318 |
| Reduced number of N (bp) | 0 | -674,896 |
| N % | **0.62%** | **0.36%** |
| Total bases without N in scaffolds (bp) | 258,891,444 | 266,296,227 |
| **Increased bases without N in scaffolds (bp)** | **0** | **7,404,783** |
| **BUSCO score (v5.2.1 embryophyta_odb10)** | **97.90%** | **98.10%** |

### Computational costs and *hardware* configuration of LOCLA

Our experiments were mostly performed on servers with a 96-core or a 160-core CPU. The detailed hardware configuration and software information are presented in Supplementary Table 10. During the experiment on LLD0021C, we analyzed the runtime of each LOCLA module on the 96-core server (Supplementary Table 11). Both LCB gap filling and LCB scaffolding require significantly more computing time than GCB gap filling or GCB scaffolding do. A closer inspection revealed that the most time-consuming process for both LCB modules was the contig assembly (Supplementary Table 12 and Supplementary Figure 9). We speculated that our barcode selection strategy could be responsible for this result; although poorly aligned barcodes were filtered through gap flanks, numerous reads belonging to the chosen barcodes were still recruited. Thus, the massive number of input reads increased with the total runtime of the SPAdes assembler. To overcome this problem, assembling tasks were run in parallel instead of sequentially. This strategy was tested with eight Microsoft Azure virtual machines (VMs; each with 72 vCPUs and 144 GB of RAM); 14 gaps were assembled simultaneously on each VM. This technique successfully reduced the total runtime to approximately 15.5 hours, improving the time efficiency substantially (Supplementary Table 13). For CHM13, all processes were run on a 160-core CPU server. Supplementary Table 14 reveals that using more CPU cores and increasing the task parallelism considerably reduces the runtime.

## Discussion

We propose a *de novo* genome assembly optimization tool that combines the advantages of state-of-the-art sequencing platforms to improve quality by enhancing genomic content. For the human samples LLD0021C and CHM13, LOCLA successfully increased the total genome size of both assemblies. The additional content improved the resolution of functionally important regions and duplicated sequences; abundant LOCLA-filled sequences were identified in protein coding regions, lncRNA, and pseudogenes. Furthermore, LOCLA identified additional repeat sequences in an analysis based on the GRCh38 human reference genome.

Despite our efforts, unresolved intra-scaffold gaps remained in the LOCLA assemblies. Some of these were too large to be crossed with a single contig, resulting in the partial filling of these regions. During LCB and GCB gap filling, only the contig with the highest alignment score was used to fill the gap to avoid excessive computational costs. Thus, the gap-filling processes could be iterated to further shorten the partially filled gaps and maximally exploit the local and global contigs. We also considered aligning both the Linked-Reads and draft assembly with the reference genome to position the barcodes more precisely (e.g., by eliminating barcodes whose reads are scattered on different chromosomes).

Moreover, functional annotation on structural variants remains vital. The linear structure of the reference genome is unable to represent the diversity of human populations ^44^. The structure of the graph genome has been developed and continues to improve. By using the graph genome ^45–48^, novel variants identified in LOCLA assemblies can be included in the genome presentation at the family or population scale. This genomic data can be further mapped to a graph reference genome and then be subjected to variant analysis with up-to-date variant callers, such as GATK^65^ and DeepVariant^66^. This procedure provides precision and personalized investigations into not only common variants but also difficult-to-detect variants. In our analysis, pervasive repetitive elements spanning approximately 45% of the genome were identified; their functional importance remains unclear. Among these, 218 repeat patterns were identified by our assembly exclusively; most of these were simple tandem repeats. Several disorders, such as neuron degenerative diseases, are known to be strongly linked to the repetitive structure of particular genomic segments ^49, 50^. Moreover, repetitive elements have been exploited and developed into genetic markers ^51–53^. On the population level, patterns of repeat sequences could be a useful link for tracing demographic changes ^54, 55^. These patterns can be used as markers or even be causal variants for disease discovery and ancestry tracing.

In spite of the discontinuation of 10xG Chromium Genome and Exome product lines, LOCLA doesn’t lose its value. There are still a myriad of genomes assembled primarily using 10xG Linked-Reads on international databases such as NCBI. LOCLA could help optimize these assemblies.

## Methods and Materials

In the following, we introduce the methods and materials used in the experiments. The datasets employed were the LLD0021C, CHM13, HG002 and *Bruguiera sexangula.* The methods included Supernova assembly, Canu assembly, Bionano Hybrid Scaffold, RagTag Scaffolding, LOCLA algorithms, and functional analysis. We also summarize our evaluation of the LOCLA assemblies on all samples.

### LLD0021C, CHM13 and HG002 data sets

#### LLD0021C data set

The LLD0021C data set comprises two subsets: Linked-reads available in NCBI SRA SRX7889242 and Bionano optical consensus maps for a sample from a Taiwanese human provided by Kwok *et al.* of the Institute of Biomedical Sciences, Academia Sinica. The 10xG Linked-Reads were sequenced on the Illumina NovaSeq 6000 instrument and yielded approximately 60x coverage of 151 bp × 151 bp paired reads (i.e., PE sequences with a length of 151 bp and sequencing depth of 60), with the total size being 191.9 Gbp. The Bionano single-molecule maps were *de novo* assembled into consensus genomic maps following the Bionano Solve Single-Enzyme Hybrid Scaffold Pipeline ^8^ using DLE-specific parameters.

#### CHM13 data set

The 10xG Linked-Reads and PacBio HiFi reads of the human sample CHM13htert cell line were obtained from the GitHub website of the T2T Consortium ^12^ in the format of raw FASTQ files. A NovaSeq instrument was used to generate 41x 150 bp × 150 bp paired 10xG reads with a total size of approximately 180 Gbp. For PacBio HiFi reads, 100 Gbp of data (32.4x coverage) in 20 kbp libraries (NCBI SRA Accession: SRX7897685-SRX7897688) and 76 Gbp of data (24.4x coverage) in 10 kbp libraries (NCBI SRA Accession: SRX5633451) were generated from PACBIO_SMRT (Sequel II) instruments.

#### HG002 data set

The PacBio HiFi reads of HG002 were obtained from the GitHub website of the Human Pangenome Reference Consortium (HPRC)^71^. SMRTbell libraries were prepared and size-selected with SageELF to the targeted length (15 kb, 19 kb, 20 kb, or 25 kb). The total size of the dataset is approximately 255 Gbp with 85.1x coverage.

### Supernova assembly and Bionano hybrid scaffolds

#### Supernova assemblies of LLD0021C and CHM13

The Supernova assembler first demultiplexes molecules and then adopts a de Bruijn graph strategy to produce an initial genome graph ^56^. Supernova also uses read pairs to cross short gaps and uses molecules in the 10x partitions to bridge gaps between two scaffolds. We used Supernova v2.0 ^10^ to produce a draft assembly of LLD0021C containing 3171 scaffolds and Supernova v2.1 to produce a draft assembly of CHM13 containing 4999 scaffolds. Both drafts were generated in the form of pseudo haploids. The commands used to run this process are provided in the Supplementary Notes.

#### Merging Bionano consensus maps and the Supernova assembly of LLD0021C into Hybrid Scaffolds

The Bionano Solve Single-Enzyme Hybrid Scaffold Pipeline, named the Bionano Pipeline for short, suggests that input scaffolds should be at least 100 kbp in length to produce high-quality hybrid scaffolds. Thus, a subset of the Supernova draft assembly (258 scaffolds were longer than 100 kbp) and the Bionano consensus map assembly were merged. A total of 68 scaffolds containing conflicting junctions were removed during this process. Conflicting junctions are loci at which the labels, marked by the Bionano enzyme DLE-1, in the two input assemblies are inconsistent. Consequently, 116 hybrid scaffolds were used. The commands used to run this process are also provided in the Supplementary Notes.

### Preprocessing of 10xG Linked-Reads by LOCLA

LOCLA comprises one preprocessing module and four main modules. The first step of LOCLA is processing of the raw reads generated by the 10x Genomics Chromium system. The preprocessing module comprises two parts:

#### Classifying and purifying FASTQs

First, Linked-Reads are classified by their barcode. Barcode information is extracted from each read and appended to the read name in the FASTQ files by using Longranger ^57^. These FASTQ files are then split into Read 1 and Read 2 (R1 and R2). Redundant bases attached to the 10xG reads are removed. Specifically, a 16 bp 10x barcode, 6 bp random primer, and 1 bp of low-accuracy sequences are trimmed from an N-mer oligo in the R1 reads, and Illumina adapter contaminants are cut from each read pair by using Trim Galore. Because polymerase chain reaction amplifies DNA fragments during Illumina sequencing, read pair duplication is inevitable ^58^. Although duplicated read pairs are known to commonly induce false positive calls in variant calling, the results of our pilot study revealed that they could also affect barcode selection and further negatively influence the method’s gap-filling performance (Supplementary Note 2). Therefore, the removal of duplicated read pairs is critical during data preprocessing to ensure that every read pair is unique. The output of the preprocessing module is a list of barcodes, each containing redundancy-trimmed and non duplicated read pairs.

#### Aligning and Filtering the read pairs

The read pairs are then mapped to the genome scaffold set using *BWA* in the end-to-end mode *BWA mem* ^59^. Secondary, duplicated, supplementary, and chimeric alignments are filtered out using *sambamba* to maintain the properly aligned read pairs^60^. Later, reads with mapping identity > 0.7 and mapping quality = 60 are retained. Mapping Quality Scores quantify the probability that a read is misplaced and are usually reported on a Phred scale67. Therefore, a Phred Score of 60 in MQ would be equivalent to an accuracy of 99.9999% in the alignment. Mapping Identity is calculated by dividing the length of sequence matches by the sum of the lengths of sequence matches, mismatches, insertions, and deletions. It demonstrates the closeness between two sequences. This filtering step ensures that the remaining read pairs are aligned with high quality, which contains convincing mapping information for subsequent work. Barcodes are then selected from this high-quality read set for use in the four main modules of LOCLA.

### LOCLA algorithms

The four main modules are LCB gap filling, GCB gap filling, LCB scaffolding, and GCB scaffolding. The basic concept of each method is presented in Fig. 2.

#### A. LCB gap filling

##### Barcode Selection

Selection of barcodes plays a crucial role in LCB gap filling. Among the three partition strategies (Supplementary Note 3), our pilot study revealed that the most effective and efficient method of assembling contigs is by selecting barcodes on the basis of each gap. First, barcodes in the high-quality read set with a sufficient number (default of three) of read pairs aligned to each scaffold are gathered. Then, barcodes with a sufficient number of read pairs (default of two) mapped within the gap flanks are collected. The flank size varies depending on the gap length as shown in the following equation:

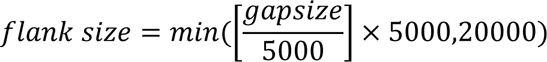

##### De novo assembling of contigs

After a barcode has been selected into the barcode list, all reads of the barcode are used to fill gaps regardless of whether the reads are in a high-quality mapped set. We use SPAdes assembler ^61^ to construct contigs. SPAdes is a genome assembly algorithm based on graph-theoretical operations on *k-*mer patterns for constructing multisized de Bruijn graphs. We denote these contigs as local contigs or “L-contigs” for short.

##### Filling gaps in scaffolds

The gap-filling algorithm comprises three steps. In the first step, the expanded mapping segments of an L-contig are identified. In the second step, each gap on a scaffold is marked as fully covered, partially covered, or unfillable simply by looking at its distance with respect to the expanded mapping segments of the L-contigs. The fully covered and partially covered gaps are then further examined using the alignment information to decide whether they can be filled by the corresponding expanded mapping segments. L-contigs with length > 1 kbp and at least 2x coverage are aligned to the target scaffold under processing by using *BWA-mem*. The alignment of an L-contig to its target scaffold usually requires more than one aligned segment. The longest putative continuing mapping range is denoted the expanded mapping segment; the remaining segments of an L-contig are trimmed on the basis of the alignment result. Beginning from the longest aligned segment with the highest alignment score, the two ends of this segment are checked and neighboring segments are iteratively merged if they are both in the right order and at a small distance. Segments far away from the main longest aligned segment are processed independently. Thus, a mapping range of the L-contig is defined; this is denoted the contig’s expanded mapping segment. For each L-contig, each gap on the target scaffold is then classified in accordance with the gap’s location with respect to the L-contig’s expanded mapping segment. If a gap is located within the segment, it is marked as fully covered. If one end of a gap is located outside the segment but within a small distance (default 50 bp), the gap is marked as partially covered. Gaps that are not covered by any contig are marked as unfillable. Some gaps can be marked as both fully covered and partially covered by different L-contig’s expanded mapping segments. These gaps are marked as fully covered. To fully fill a gap with an expanded mapping segment, the mapping identity must be greater than 80 on both flanks of the gap. For partially covered gaps, the gap is filled on each flank by using the expanded mapping segment with the highest score among those segments with more than 300 matched base pairs and mapping identity > 90. Note that a gap with both flanks comprising duplicated sequences is not considered a candidate for gap filling.

#### B. GCB Gap Filling

This module is an alternative for LCB gap filling. Instead of being filled with *de novo* assembling contigs, gaps are filled with global contigs (G-contigs), which serve the sole purpose of filling gaps in earlier assemblies. G-contigs are foreign TGS long reads or foreign scaffolds produced by the sequence assembler. Initially, gaps of foreign long reads or scaffolds that are longer than a specific length (default 20 bp) are detected, and the G-contigs are then broken into smaller fragments. These G-contig fragments are then aligned with the entire assembly by using *minimap2* and used to fill gaps with the same algorithm as for LCB gap filling.

#### C. LCB Scaffolding

##### Defining candidate scaffold pairs from the barcode distribution on scaffolds

On the basis of the filtered read-to-scaffold alignment obtained in the preprocessing module, the algorithm tallies and records all barcodes of the reads that were mapped onto the head and tail of each scaffold (default of 20 kbp). For each scaffold end, the two scaffold ends of other scaffolds that share the largest number of barcodes with it are kept. This list of scaffold end pairs is denoted the list of candidate scaffold pairs (CSPs).

##### De novo assembly of L-contigs

All reads with the same barcodes are used to construct contigs for each CSP by using the SPAdes assembler.

##### Concatenating scaffolds with high-quality contigs

L-contigs are aligned to the CSP with *BWA-mem*, and those L-contigs with mapped length > 1000 bp and mapping identity > 0.7 within 20 kbp of both CSP ends are kept. Finally, each CSP is connected with the L-contig with the highest mapping identity.

#### D. GCB Scaffolding

The algorithm is fundamentally the same as that for LCB scaffolding but with a few alterations. First, instead of L-contigs, G-contigs are used as the input. Second, the alignment tool is *minimap2* rather than *BWA-mem*.

### Functional analysis of LOCLA

Useful genes as well as repeated elements were discovered in the additional genomic content identified by LOCLA.

#### Gene Annotation

Genome assemblies were aligned to GRCh38 and CHM13 reference genomes to compare their genomic content and the sizes of gap regions. We employed *minimap2* to perform alignment and identify the increased genome content with direct comparisons based on the two reference coordinates. The increased genome content was then annotated on the basis of GENCODE v29 ^16^.

#### Repeat Element Identification

To evaluate the performance of different assembly pipelines in the repeat regions, RepeatMasker was applied to identify the repeat elements and their corresponding repeat patterns. RepeatMasker was run with the *species human* option in settings. The outputs of RepeatMasker were further processed using the Perl script *onecodetofindthemall.pl* to categorize the repeat elements into several repeat patterns and generate copy number estimates^62^.

#### Gene Prediction for unmapped scaffolds

To investigate whether the unmapped scaffolds actually existed in the genome, we applied AUGUSTUS ^17^ for gene predictions. We assumed that if inferred genes were located in these scaffolds, these scaffolds may exist in the genome. Augustus was run with the *--species=human--UTR=on* settings. To further validate these inferred genes, we performed protein BLAST ^18^ to determine whether the predicted sequences were conserved in organisms.

### Summary of the LOCLA assembly of the three human datasets

The entire workflow for LLD0021C is presented in Figure 3a. The initial draft containing 3,171 scaffolds was generated by Supernova v2.0. We then performed LCB gap filling on 1171 scaffolds longer than 22 kbp. A subset of 258 gap-filled scaffolds with length > 100 kbp were then merged with the optical consensus maps of Bionano Genomics to create 116 hybrid scaffolds. After retrieving the 68 scaffolds unused by Bionano Solve and the 2913 scaffolds shorter than 100 kbp, we applied the LOCLA modules to the whole assembly in the following order: GCB gap filling, LCB gap filling, LCB scaffolding and finally GCB Scaffolding. For CHM13, a draft assembly containing 4,999 scaffolds was produced by Supernova v2.1. Using 10xG Linked-Reads, we applied GCB gap filling, LCB gap filling and LCB scaffolding to the Supernova draft (Figure 3b). Lastly, we performed GCB Gap Filling and GCB Scaffolding with PacBio HiFi reads. For HG002 (Figure 3c), We assembled the PacBio HiFi reads by Canu^15^ first, then employed a reference-guided scaffolding tool RagTag^68^ using CHM13v1.1^12^ as reference and the Canu-assembled contigs as query. We finalize this workflow by filling gaps using the Canu-assembled contigs discarded by RagTag. Detailed descriptions of both workflows are presented in the Supplementary Notes.

**Figure 3.**
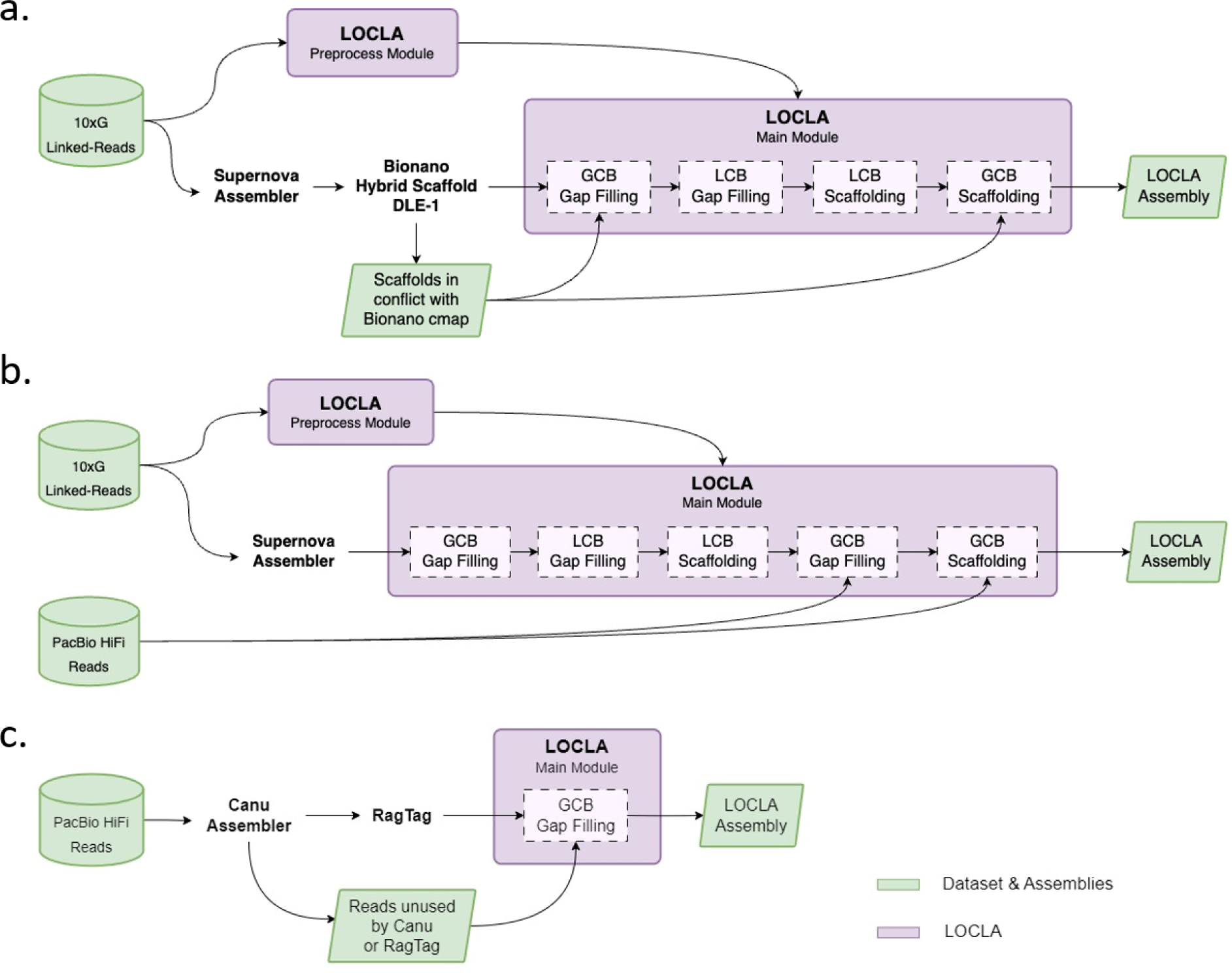
| The flexibility of LOCLA is demonstrated through different experiment pipelines. **a,** The entire pipeline of LLD0021C. Initially, we generated a pseudo haploid assembly with the Supernova assembler (v2.0). Bionano Optical Mapping (OM) only takes scaffolds over 100kbp as input, thus, we fill in gaps on a subset of scaffolds longer than the L99 length. Afterwards, we obtained a hybrid assembly and a subset of unused scaffolds: those that were in conflict with the Bionano OM cmap or shorter than 100kbp. Next, we conducted GCB Gap Filling on the hybrid assembly with the unused scaffolds. Subsequently, we merged the filled hybrid assembly and unused scaffolds into one. Next, LCB Gap Filling and LCB Scaffolding were then performed on our assembly. Finally, we conducted GCB Gap Filling to reach our final assembly. **b,** The pipeline of CHM13. We first produced a pseudo haploid assembly using the Supernova assembler v2.1. Then, we performed LOCLA modules in the order of GCB Gap Filling, LCB Gap Filling, LCB Scaffolding and GCB Scaffolding. Besides Supernova, we didn’t adopt any other existing assembling tools for the purpose of validating the performance of LOCLA. **c,** The pipeline of HG002. Initially, we utilized Canu to compile the PacBio HiFi reads. After that, we utilized a reference-guided scaffolding tool called RagTag to align the Canu-assembled contigs with CHM13v1.112 as the reference. Finally, we used LOCLA to fill in any gaps, using the PacBio HiFi reads that were not included in the Canu assembly.

### Evaluation on the three human datasets

For LLD0021C, we compared our assembly to the reference genome GRCh38.p13. For CHM13, we took the version of CHM13 v1.1 as our reference. As for HG002, the HPRC maternal haploid of HG002 was chosen as our benchmark, on account that it comprises the same chromosomes (chr1-22 and chrX) as our reference used in RagTag^68^. For global alignment, the tool used for mapping whole genome assemblies onto the reference genomes is *minimap2* ^20^. One of the presets for full-genome alignment, *asm5*, is employed here for sequences with divergence < 1%. As for local alignment, BLAST is used for comparison on subject and query sequences. For both global and local alignment, we employed the Longest Common Sequence algorithm and calculated their Mapping Identities by dividing the length of sequence matches by the sum of the lengths of the query and deletions. The tool for generating alignment plots is D-GENIES^72^ (Dot plot large Genomes in an Interactive, Efficient and Simple way), which is an online tool designed to compare two genomes.

### *Bruguiera sexangula* genome dataset and optimization of assembly via LOCLA

Illumina sequencing data from the 10× Genomics library and RNA-seq data (MGISEQ) were submitted to the NCBI Sequence Read Archive (SRA) database under BioProject accession number PRJNA734123 (DNA short-read data: SRX12279148; RNA-seq data: SRX12119193). The *Bruguiera sexangula* genome assembly published by Pootakham W. *et al.*^69^ was deposited in the DDBJ/ENA/GenBank database under the accession number JAHLGP000000000. It was constructed with RagTag^68^, taking the Supernova assembly of *Bruguiera sexangula* genome as a query and *Bruguiera parviflora* genome as reference. We then performed LCB Gap Filling and LCB Scaffolding on this draft genome.

## Code Availability

A docker image of LOCLA is freely available on our DockerHub page (https://hub.docker.com/r/lsbnb/locla), and the source code is accessible on our GitHub page (https://github.com/lsbnb/locla).

## Acknowledgements

This manuscript was edited by Wallace Academic Editing.

## Data Citation

1. Pui Kwok’s lab, IBMS SINICA. (2020). LLD0021C. https://www.ncbi.nlm.nih.gov/bioproject/626976 ; SRX7889242: 10xG Linked Reads of LLD0021C: https://www.ncbi.nlm.nih.gov/sra?LinkName=biosample_sra&from_uid=14178705

## Notes

### Competing Interest Statement

The authors have declared no competing interest.

