## Supplementary Information for "A Novel Genome Optimization Tool for Chromosome-Level Assembly across Diverse Sequencing Techniques"

1. **Supplementary Notes**

Supplementary Note 1. List of commands for LLD0021C and CHM13

Supplementary Note 2. Effects of duplicate read pairs on gap filling

Supplementary Note 3. Testing different barcode selection strategies

Supplementary Note 4. Pseudocode of LCB Gap Filling

Supplementary Note 5. Pseudocode of LCB Scaffolding

Supplementary Note 6. Assembly workflow of LLD0021C and CHM13

Supplementary Note 7. Workflow of Eel genome assembly and BUSCO evaluation

2. **Supplementary Figures**

Supplementary Figure 1. Workflow for preprocessing FASTQs

Supplementary Figure 2. Gap filling experiments on 4 scaffolds under different BS affinity with duplicate and non-duplicate reads set

Supplementary Figure 3. Three different barcode selection strategies

Supplementary Figure 4. Illustration of the basic concept of the four main LOCLA modules

Supplementary Figure 5. Workflow of filling gaps in scaffolds

Supplementary Figure 6. Evaluating the accuracy of HG002 assemblies based on alignments to their references

Supplementary Figure 7. The comparison between GUA assembly and GMA assembly

Supplementary Figure 8. Time consumption of each process in LCB Gap Filling and LCB Scaffolding on sample LLD0021C

3. **Supplementary Tables**

Supplementary Table 1. Analysis of barcoded reads against draft assembly after removing duplicate and barcodes with no more than 3 read pairs

Supplementary Table 2. Scaffold-based gap filling analysis on scaffold 21, 28, 36 and 39 using duplicate read pairs

Supplementary Table 3. Scaffold-based gap filling analysis on scaffold 21, 28, 36 and 39 using non-duplicate read pairs

Supplementary Table 4. Gap filling analysis for each partition algorithm

Supplementary Table 5. Analysis of filled gaps by LOCLA on LLD0021C

Supplementary Table 6. LOCLA outperforms Supernova in repeat sequences (sample: LLD0021C)

Supplementary Table 7. Analysis of filled gaps by LOCLA on CHM13

Supplementary Table 8. Gap closure results employing SOAPdenovo (GapCloser) and GapFiller obtained on Homo sapiens (chromosome 14)

Supplementary Table 9. LOCLA reduces a considerable amount of gaps through several iterations

Supplementary Table 10. Hardware configuration and software information

Supplementary Table 11. The runtime of each LOCLA module on the 96-core CPU server (sample: LLD0021C)

Supplementary Table 12. The runtime of LCB Gap Filling and LCB Scaffolding on the 96-core CPU server (sample: LLD0021C)

Supplementary Table 13. Reducing runtime by distributing assembling tasks on 8 Virtual Machines (sample: LLD0021C)

Supplementary Table 14. Runtime of assembling contigs (LCB Gap Filling) for LLD0021C and CHM13

Supplementary Table 15. LOCLA increased the content of disease-related genes on the Supernova assembly (sample: LLD00021C)

Supplementary Table 16. Genes revealed after enhancing Supernova by the LOCLA assembly (sample: CHM13)

1. **Supplementary Notes**

**Supplementary Note 1. List of commands for LLD0021C and CHM13.**

***Supernova for LLD0021C***

supernova run \

--id=LLD0021C_10x_supernova2 \

--fastqs=/path/to/10x_fastq/LLD0021C \

--localcores=64 \

--localmem=1024

supernova mkoutput \

--asmdir=LLD0021C_10x_supernova2/outs/assembly \

--outprefix=human_LLD0021C_10k_pseudohap \

--style=pseudohap \

--minsize=10000

***Supernova for CHM13***

supernova run \

--id=CHM13_supernova2 \

--fastqs=/path/to/CHM13/10x_fastq \

--localcores=64 \

--localmem=1024

supernova mkoutput \

--asmdir=CHM13_supernova2/outs/assembly \

--outprefix=CHM13_10k_pseudohap \

--style=pseudohap \

--minsize=10000

***Bionano Solve Single-Enzyme Hybrid Scaffold for LLD0021C***

perl hybridScaffold.pl \

-n human_LLD0021C_10k_pseudohap.fasta \

-b LLD0021C.exp_refineFinal1_merged_q.cmap \

-c hybridScaffold_config.xml \

-r /path/to/bionano/Solve3.4_06042019a/RefAligner/8949.9232rel/RefAligner \

-o IBMS_LLD0021Cpseudohap \

-f \

-B 2 \

-N 2

**Supplementary Note 2. Effect of duplicated read pairs on gap filling**

During Illumina sequencing, PCR amplifies DNA fragments, resulting in read pair duplications. Although duplicated read pairs do not typically affect other gap-filling techniques, this note demonstrates that they can affect barcode selection and thus affect gap-filling performance.

Four scaffolds—scaffolds 21, 28, 36, and 39, with lengths 5.2, 4.8, 4.3, and 4.2 Mbp, respectively—were selected from a subset of Eel Genome 1.0 [ncbi accession number] (948 scaffolds with a total size of 991.45 Mbp containing 4,372,491 bp of gaps) and used as the data set. We conducted two gap-filling experiments on these scaffolds using duplicated and nonduplicated read pairs, respectively. By aligning all read pairs to the four scaffolds, duplicated read pairs could be identified and the shorter read pairs could be discarded, resulting in only one unique read pair remaining. The statistics of the dataset after the removal of duplicated read pairs for gap filling are presented in Supplementary Table 1.

Before gap filling, a unique pair was defined as a read pair of a barcode with both reads mapped to the same scaffold and only to this scaffold. The number of unique pairs was used as the criteria for barcode selection; that is, given a target scaffold, barcodes with a sufficiently large number of unique pairs to fill its gaps were selected. Four criteria for recruiting a barcode for gap filling in a specific scaffold were tested. In each criterion, a threshold value for the number of unique pairs coexisting on the scaffold and barcode was specified: 1, 3, 5, or 10. The results of using duplicated and nonduplicated read pairs are presented in Supplementary Table 2 and 3. A comparison of the two tables reveals that in general, the percentage of the length of filled gaps increased and the computational time decreased when the duplication threshold was decreased. Thus, the removal of duplicated read pairs did improve the gap-filling performance. Supplementary Table 3 reveals that although the highest filled-gap-length percentage for scaffold 21, 28, and 36 was obtained for a threshold of 1, the runtime of the task was excessive. To optimize both the runtime and the length of filled gaps, the default threshold value for the number of unique pairs was set to 3. For scaffold 28, changing the threshold from 1 to 3 decreased the filled-gap-length percentage from 79.9% to 70.6%; however, the computational time was halved, decreasing from 62.3 to 31.6 hours.

**Supplementary Note 3. Testing different barcode selection strategies**

Three barcode selection strategies were designed: scaffold-based, gap-based, and window-based (Supplementary Figure 3). In Supplementary Note 2, barcodes are described as being recruited for each scaffold; this is a scaffold-based strategy. In the gap-based strategy, barcodes are selected by aligning them with the two flankings of each gap. In the window-based strategy, gaps located within a window (60 kbp) are grouped, and barcodes are collected on their extended flanking (5 kbp). In both the gap-based and window-based strategies, barcodes with at least 3 unique pairs aligned with the target scaffold and 2 unique pairs aligned with both flankings are recruited.

Gap filling was then performed on the pilot data (scaffolds 21, 28, 36, and 39) by using the three strategies. All experiments were performed on a server with 96 CPUs and 1.48 TB of RAM. Supplementary Table 4 presents a comparison of the gap-filling results. The gap-based and window-based strategies both significantly outperformed the scaffold-based strategy in terms of computational time and the length of the filled gaps. Although the window-based strategy required a shorter runtime, the gap-based strategy maximized the length of the filled gaps. However, the runtime of the gap- and window-based strategies differed only marginally; thus, the gap-based strategy was employed for barcode selection in GABOLA.

**Supplementary Note 4. Pseudocode of LCB Gap Filling**

**STEP I. Barcode Selection**

C70M60 sam file: The alignment of 10x linked reads to our draft assembly with mapping identity greater than or equal to 70 and mapping quality greater than or equal to 60.

{BXlist}S: Barcode list for each scaffold.

{BXlist}S.G: Barcode list for each gap per scaffold.

```
PROGRAM ProduceBXList (C70M60 sam file, draft assembly):
 Rename scaffolds by descending order of size;
 FOR each scaffold DO
  IF (number of mapped read pairs >= 3)THEN
```

   Add barcode to {BXlist}S;

```
  ENDIF
 ENDFOR
 FOR each gap on scaffold DO
  Determine flanking region based on gap size;
  IF (number of mapped read pairs on gap’s flanking >= 2)THEN
```

Select barcode from {BXlist}S and add barcode to {BXlist}S.G;

```
ENDIF
 ENDFOR
```

**IF** (number of barcodes in {BXlist}S.G <= 10)**THEN**

```
Gap will not be processed;
 ENDIF
END

STEP II. De novo Local Assembly of Contigs
Nondup fastq: Fastq file of non-duplicated 10x reads split according to barcode type.

PROGRAM Assemble (nondup fastq):
 Determine number of gaps to process simultaneously;
 FOR each gap
```

**DO** Collect barcoded reads based on {BXlist}S.G from nondup fastq;

```
Assemble reads into contigs;
 ENDFOR
END

STEP III. Gap Filling
Lc: Left-most position of the longest contig mapping range on a scaffold.
Rc: Right-most position of the longest contig mapping range on a scaffold.
I: Set of longest contig mapping intervals.
MI: Mapping identity of contig on scaffold.
ML: Length of matched base of contig on scaffold.
U: Set of unmapped segments of contig on scaffold.

PROGRAM Fill (draft assembly):
 FOR each contig DO
  IF (contig length >= 1kbp  contig coverage >= 2)THEN
Align contig to scaffold;
IF (contig is not within the flankings of any gap) THEN
     Remove contig;
   ELSE
     {Lc, Rc} = longest mapping range of contig on the scaffold;
     I = I  {Lc, Rc};
   ENDIF
  ENDIF
 ENDFOR
 FOR each gap DO
IF (gap  {Lc, Rc}) THEN
Gap is considered fully-covered;
FOR each contig covering the gap DO
WHILE ( MI >=80 on both flankings of gap) DO
 IF ( U is only within the gap) THEN
  Gap is fully filled;
 ELSE
  Gap will not be filled;
 ENDIF
ENDWHILE
    ENDFOR
ELIF (gap  {Lc-50bp ,Rc}  {Lc, Rc+50bp}) THEN
Gap is considered partially-covered;
FOR each contig partially-covering the gap DO
WHILE ( MI >=90 on the flanking of gap  ML >=300bp ) DO
 Fill part of the gap;
  ENDWHILE
ENDFOR
ELSE
Gap is considered unfillable;
ENDIF
 ENDFOR
END
```

**Supplementary Note 5. Pseudocode of LCB Scaffolding**

**STEP I. Determine candidate scaffold pairs**

*E*: Candidate of scaffold end pairs.

*L*: Barcode list on each scaffold head and tail.

*F*: Filtered candidate of scaffold end pairs.

**PROGRAM** Preprocessing (*E,L*):

**FOR** each scaffold pair in *E* **DO**

ScafA, ScafB = scaffold end pair

**IF** (ScafA==ScafB) **THEN**

Remove scaffold pair;

**ELIF** (ScafA or ScafB appear more than two times in *E*) **THEN**

Add only the scaffold pairs with first and second most

barcode counts to *F* ;

**ENDIF**

**ENDFOR**

**FOR** each scaffold pair in *F* **DO**

ScafAf , ScafBf = scaffold end pair;

Collect barcode list on ScafAf end from *L*;

Collect barcode list on ScafBf end from *L*;

**ENDFOR**

**END**

**STEP II. *De novo* Assembling of Contigs**

*F*: Filtered candidate of scaffold end pairs.

Nondup fastq: Fastq file of non-duplicated 10x reads splited according to barcode type.

**PROGRAM** Assemble (*F*, nondup fastq):

Determine number of scaffold pairs to process simultaneously;

**FOR** each scaffold pair in *F* **DO**

ScafAf , ScafBf = scaffold end pair;

Collect reads from nondup fastq based on barcode list on ScafAf;

Collect reads from nondup fastq based on barcode list on ScafBf;

Assemble reads into contigs;

**ENDFOR**

**END**

**STEP III. Scaffolding**

*F*: Filtered candidate of scaffold end pairs.

Cf: Candidate contigs for each scaffold pair in *F*

*MI*: Mapping identity of contig on scaffold.

*ML*: Length of matched base of contig on scaffold.

**PROGRAM** Scaffolding (*F*, draft assembly):

**FOR** each scaffold pair in *F* **DO**

Align Cf to draft assembly;

**FOR** each contig in Cf **DO**

```
WHILE (contig is aligned within 20kbp of both scaffold ends)
DO
IF ( ML >= 1kbp and MI >= 70) DO
Concatenate two scaffolds;
ENDIF
ENDWHILE
ENDFOR
 ENDFOR
END
```

**Supplementary Note 6. Assembly workflow of LLD0021C and CHM13**

***Pipeline of LLD0021C assembly.*** The experiment had two stages. First, the Supernova assembler was run, and LCB gap filling and Bionano Solve were performed. Subsequently, GABOLA was applied. By running Supernova v2.0^19^ on the 10x Linked-Reads, a haplotype draft assembly of 3171 scaffolds was generated. The gaps were then filled with LCB Gap Filling on a subset of the assembly containing 1171 scaffolds longer than the N99 length (22,459 bp) before the Bionano Pipeline was implemented. The Bionano Pipeline only accepts scaffolds with lengths greater than 100 kbp as its input; therefore, sequences shorter than 100kbp or in conflict with the Bionano cmap were set aside during the process; 258 scaffolds remained for the next stage. The output of Bionano Solve comprised a set of 116 Hybrid Scaffolds.

In the second stage of the pipeline, GCB gap filling was performed on the 116 hybrid scaffolds with the 2981 unused sequences. LCB gap filling was next conducted on the assembly through the gap-based strategy. Finally, GABOLA LCB and GCB scaffolding were applied; 244 scaffolds were connected, yielding a final draft assembly of 2975 scaffolds.

***Pipeline of CHM13 assembly.*** Similar to the final workflow for LLD0021C, the 10x Genomics Software Supernova v2.1^23^ using Linked-Reads was used to generate a haplotype draft assembly of 4999 scaffolds. However, Bionano OM was not included in this pipeline. The same procedure as for the second stage of LLD0021C was performed; GCB gap filling was first used to close the largest gaps. LCB gap filling was next applied to the assembly. Finally, 212 scaffolds were connected with LCB and GCB Scaffolding, resulting in a final draft assembly of 4809 scaffolds.

**Supplementary Note 7. Workflow of eel genome assembly and BUSCO evaluation *Constructing the eel genome.*** The data set comprised 200x coverage Illumina reads [both PE and five MPs (2, 4, 8, 10, and 15 kb)], 8x coverage PacBio long reads, and 60x coverage 10xG Linked-Reads. The PE and MP reads mapped with low quality were first filtered using subset selection methods and a sequencing quality assessment tool, SQUAT. The initial draft assembly (Eel genome 1.0) was then generated using ALLPATHS-LG^32^, SSPACE^33^, GapCloser^34,35^, and Canu^36^. Multiple processes were then performed on the draft, including SALSA^37^, SSPACE-long^38^, and ALLMAPS^39^. We also conducted several iterations of GCB gap filling with PacBio long reads as the input; each time, used contigs from the previous round were filtered out. Subsequently, 10xG Linked-Reads were used to perform LCB gap filling and LCB scaffolding. Finally, we used POLCA, a polishing tool for error correction, to achieve the final version of the assembly.

***BUSCO evaluation for eel genome.*** BUSCO (the Benchmarking Universal Single-Copy Ortholog assessment tool) evaluates the quality of a genome assembly by using predefined gene models of single-copy orthologs; the tool selects orthologous groups of genes present in at least 90% of species including arthropods, vertebrates, metazoans, fungi, eukaryotes, and bacteria. The completeness of a genome can be estimated by searching for these genes in the genome draft, which is scanned by using tblastx^41^, hidden Markov models, and AUGUSTUS^42^. The predicted genes are marked as “complete” or “duplicate” if the gene models in the BUSCO algorithm are satisfied. Some gene model matches are partial and are marked as fragmented; the remaining BUSCO groups with no matches are marked as missing. BUSCO was used to evaluate the genomic content of our assembly and determine whether the GABOLA gap-filling pipeline had identified additional gene content. For the quality assessment, we used BUSCO v3.0.2 in “genome” mode and specified “zebrafish” as the reference species and “Vertebrata” as the taxonomy.

2. **Supplementary Figures**

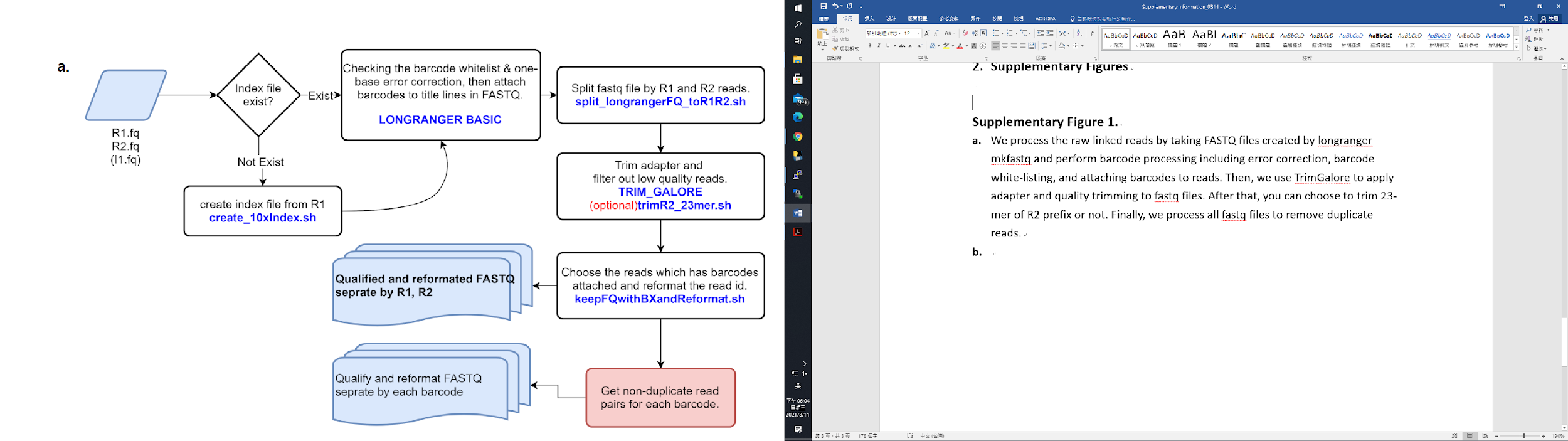

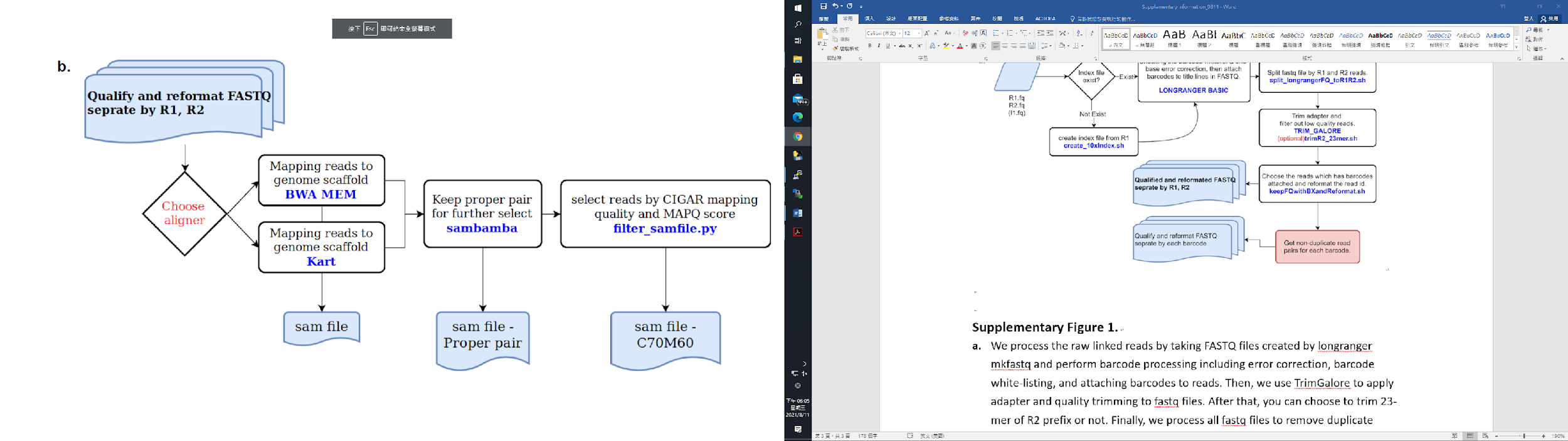

**Supplementary Figure 1. Workflow for preprocessing FASTQs**

a. We process the raw linked reads by taking FASTQ files created by *longranger* *mkfastq* and perform barcode processing including error correction, barcode white-listing, and attaching barcodes to reads. Then, we use *TrimGalore* to apply adapter and quality trimming to fastq files. After that, you can choose to trim the 23-mer of R2 prefix or not. Finally, we process all fastq files to remove duplicate reads.
b. We map reads to the genome and filter them: We begin with aligning reads to genome scaffolds by *BWA-mem* or *Kart*. Then, we keep proper pairs by *SAMBAMBA (v0.6.9)*. Finally, we filter the alignment by match quality calculated from CIGAR string and MAPQ score.

#definition of proper pair: mapped and not {secondary_alignment or duplicate or supplementary or chimeric}

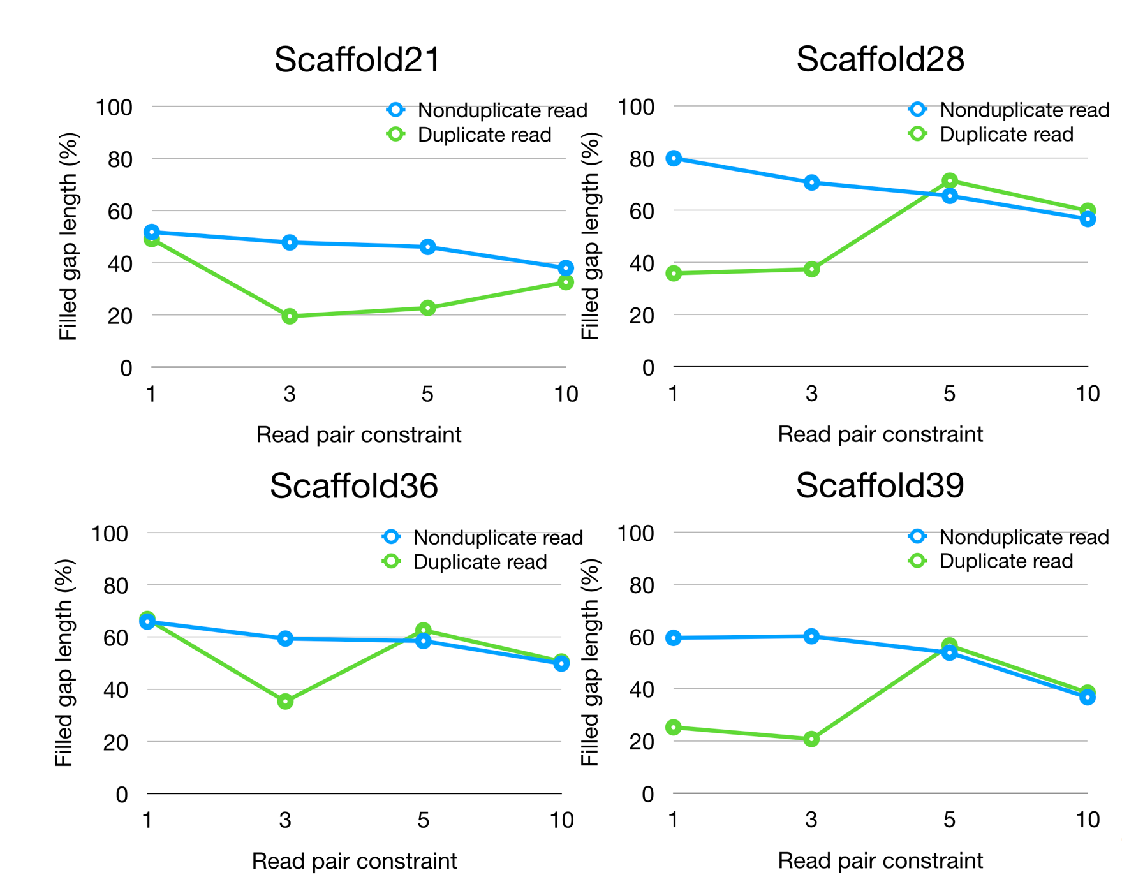

**Supplementary Figure 2. Gap filling experiments on 4 scaffolds under different BS affinity with duplicate and non-duplicate reads set**

The four line plots suggest that gap filling with non-duplicate reads performs better than duplicate reads when the read pair constraint is set as 1 or 3. Non-duplicate reads deliver exceeding results especially under when the constraint is 3 read pairs. All four scaffolds filled in at least 20% more gaps, with the maximum number of 40% on scaffold 39. However, the effect of utilizing non-duplicate reads decreases with the increase of read pair constraint number.

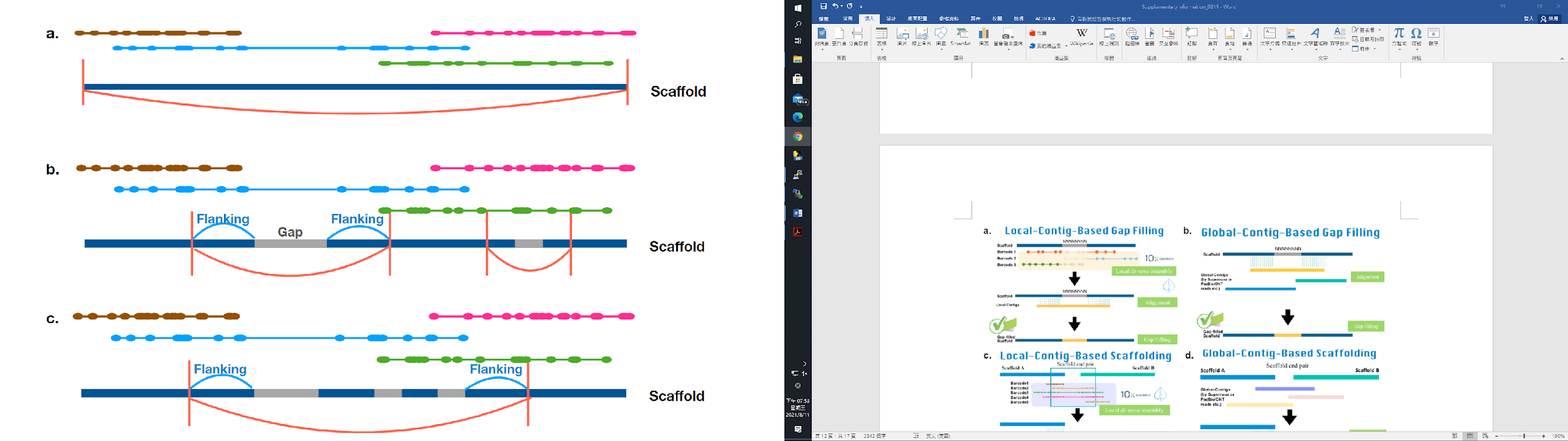

**Supplementary Figure 3. Three different barcode selection strategies**

a. Scaffold-based barcode selection: We collect barcodes mapped onto the entire scaffold
b. Gap-based barcode selection: We collect barcodes mapped onto the flanking region of each gap. The flanking size is determined by the gap size.
c. Window-based barcode selection: We group gaps located within a window of 60kbp and collect barcodes mapped onto the flanking region (5 kbp) per window.

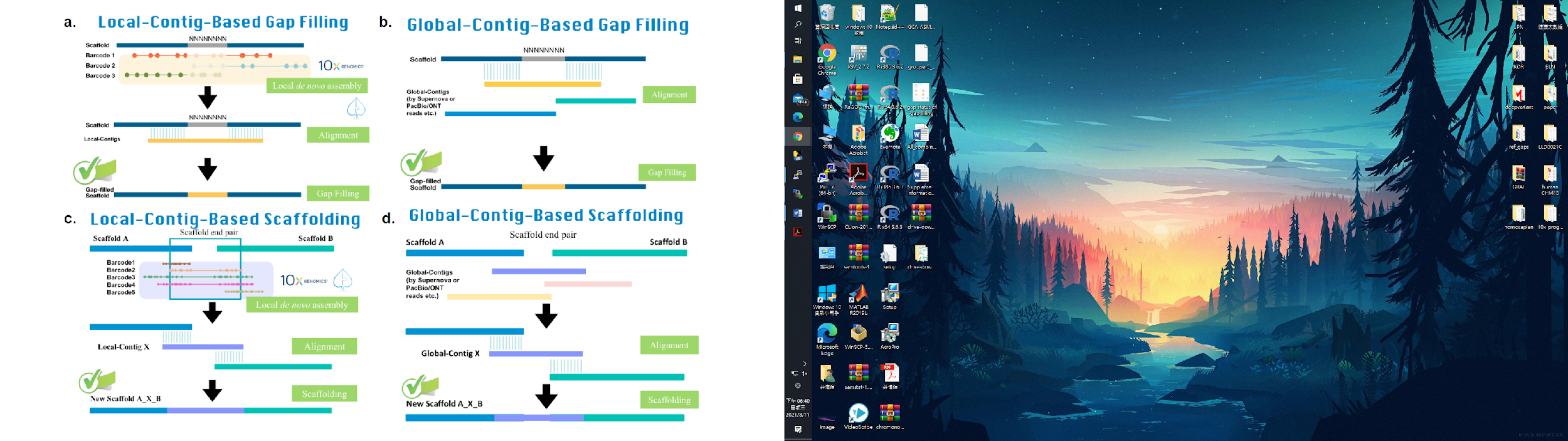

**Supplementary Figure 4. Illustration for the basic concept of the four main LOCLA modules**

a. “Local-Contig-Based (LCB) Gap Filling”: First, we align all barcoded linked-reads to the scaffolds and *de novo* assemble Local-contigs using the reads belonging to barcodes mapped within gap flankings. Then, we map these contigs onto the scaffolds and determine the best hit to fill in gaps.
b. “Global-Contig-Based (GCB) Gap Filling”: We align Global-contigs to all scaffolds and find the best hit to fill in gaps.
c. “Local-Contig-Based (LCB) Scaffolding”: We align all barcoded linked-reads to the head and tail of scaffolds and pair up scaffolds with shared barcodes. Then we construct Local-contigs from the barcoded-reads and connect scaffolds with the optimal L-contig.
d. “Global-Contig-Based (GCB) Scaffolding”: Identical to LCB Scaffolding, we align linked-reads to the ends of scaffolds and pair up scaffolds with shared barcodes. Global-contigs are then mapped onto all scaffolds pairs and connected with the most ideal G-contig.

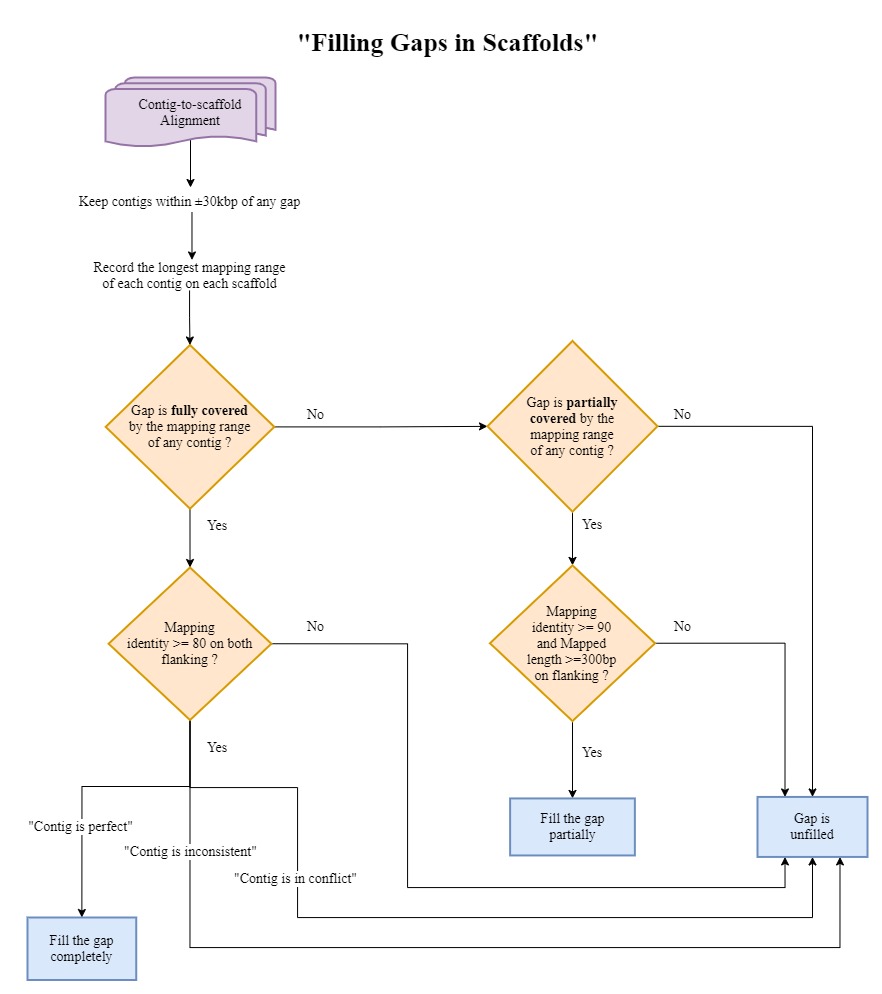

**Supplementary Figure 5. Workflow of filling gaps in scaffolds**

First, gaps are classified into three types based on contig-gap flanking alignments. For those fully-covered gaps, we check the mapping quality of the extended contig segment to the gap flanking, and see whether it meets the criteria in both flanking to replace the gap regions. To avoid introducing the misassembled contig into the previous assembled genome, only the perfect contigs are capable of filling gaps.

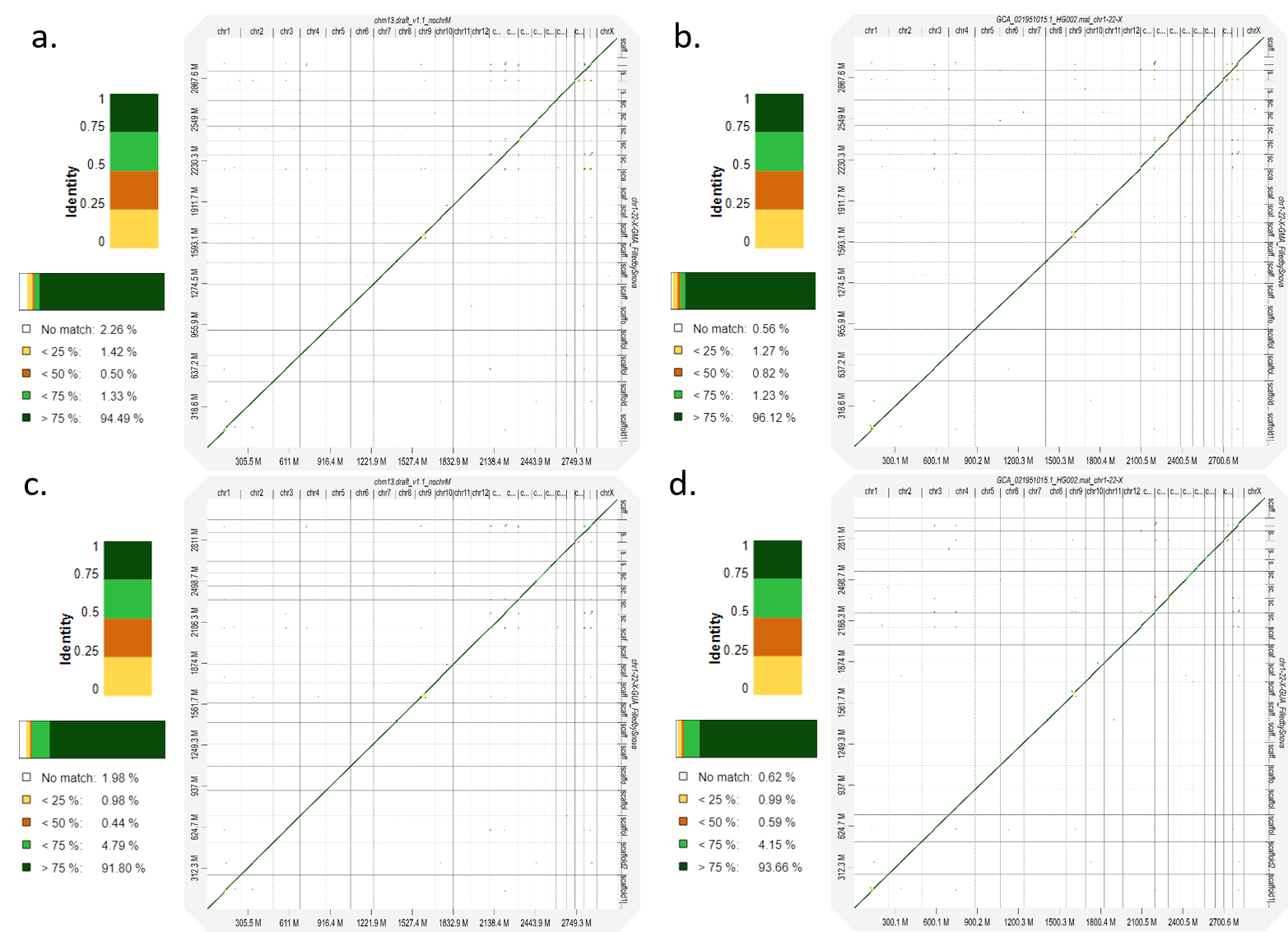

**Supplementary Figure 6.** **Evaluating the accuracy of HG002 assemblies based on alignments to the references.**

a, The alignment between the gap-filled GMA assembly and the CHM13 v1.1 assembly (chr1~22, X). b, The alignment between the gap-filled GMA assembly and the HPRC maternal haploid assembly. c, The alignment between the gap-filled GUA assembly and the CHM13 v1.1 assembly (chr1~22, X). d, The alignment between the gap-filled GUA assembly and the HPRC maternal haploid assembly.

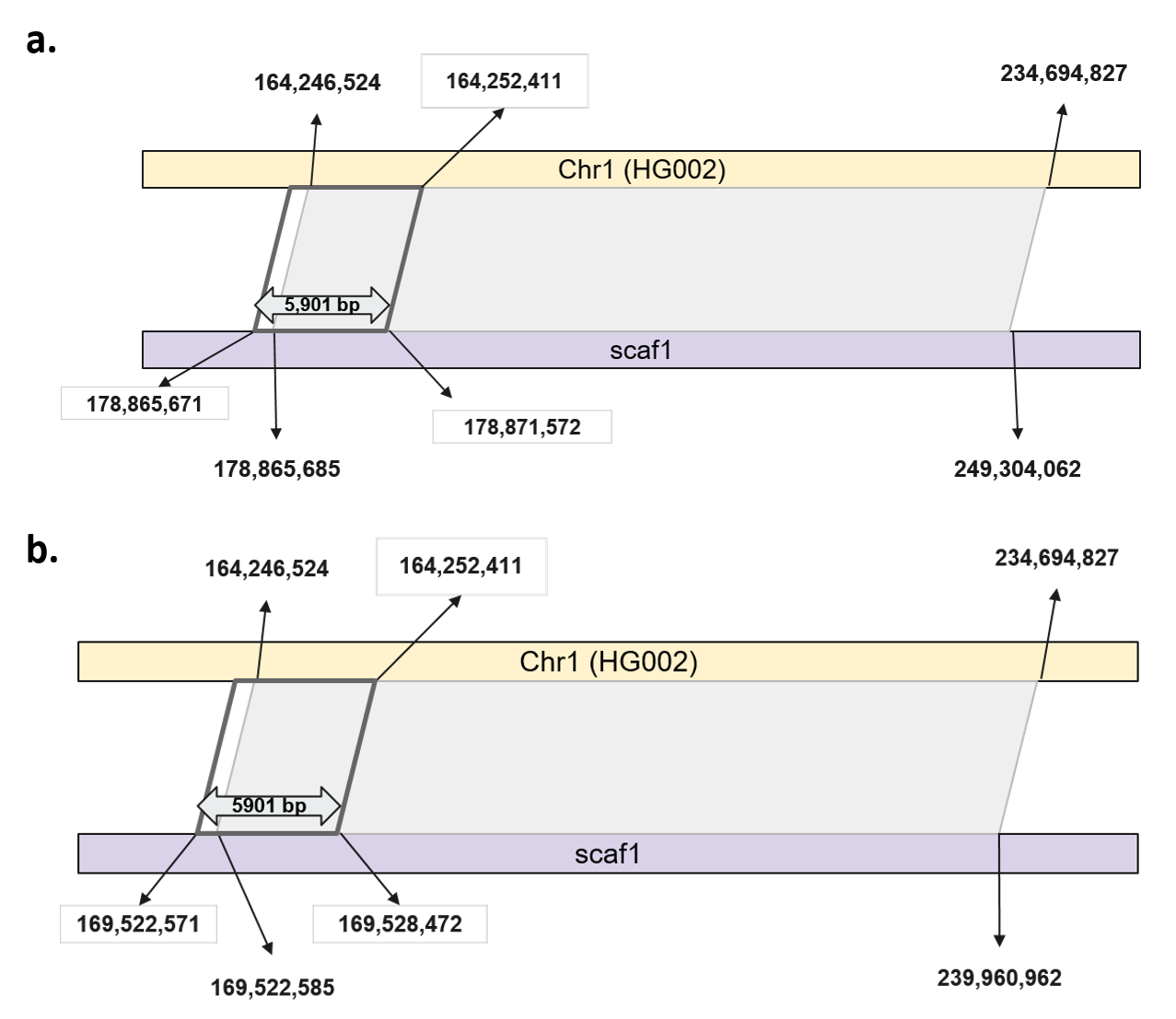

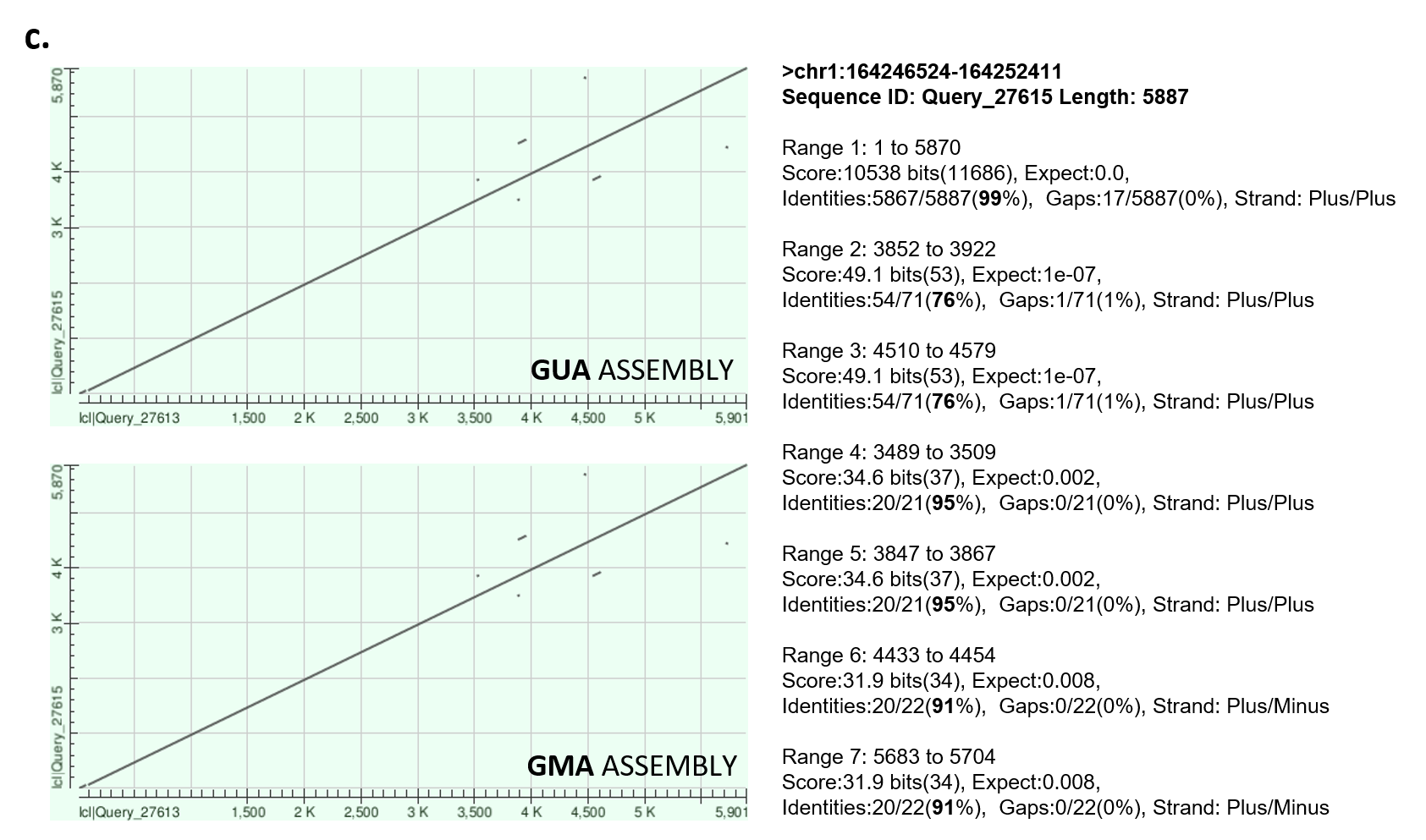

**Supplementary Figure 7. Global alignment of scaffold 1 on chromosome 1 of GUA and GMA assemblies**

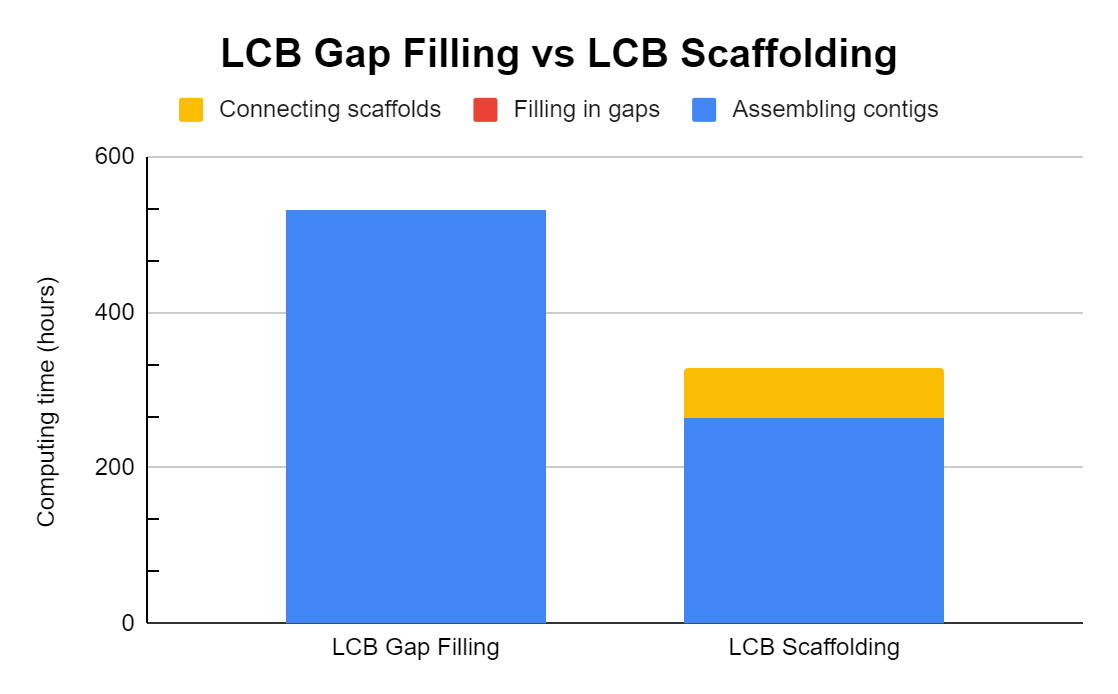

**Supplementary Figure 8. Time consumption of each process in LCB Gap Filling and LCB Scaffolding on sample LLD0021C**

We recorded the runtime of each stage in the two modules on a 96-core CPU server: (1) “Assembling contigs” and “Filling in gaps” for LCB Gap Filing, respectively around 531 hours and 1.5 hours. (2) “Assembling contigs” and “Connecting scaffolds” for LCB Scaffolding, respectively about 265 hours and 64 hours. The process of contig assembly takes up most of the time in both modules.

3. **Supplementary Tables**

**Supplementary Table 1. Analysis of barcoded reads against draft assembly after removing duplicate and barcodes with no more than 3 read pairs**

|  | **# of Reads** | **# of Barcodes** | **#Scaffold with at least**  **one mapped read** |
| --- | --- | --- | --- |
| Barcoded read | 1,177,151,368 | 3,449,903 | - |
| Trimmed read | 1,142,645,930 | 3,423,181 | - |
| Non-duplicate read | 931,300,700 | 2,395,977 | - |
| Aligned read | 931,300,700 (100%) | 2,395,977 (100%) | 7,804 (100%) |
| Proper pair alignment | 652,825,738 (70.1%) | 2,262,747 (94.4%) | 7,801 (99.9%) |
| Mapping identity > 70 and  Mapping quality = 60 | 280,540,246 (30.1%) | 1,772,060 (74.0%) | 6,126 (78.5%) |

**Supplementary Table 2. Scaffold-based gap filling analysis on scaffold 21, 28, 36 and 39 using duplicate read pairs**

|  | **# of unique pairs** | **#Recruited Barcode** | **#Recruited Read Pairs** | **Wall Clock Time (hr)** | **Filled Gap Length (%)** |
| --- | --- | --- | --- | --- | --- |
| **Scaffold21**   - **5.2Mbps,** - # Gaps: 313 - **# N: 383.3Kbps** | 1 | 178,719 | 69,002,194 | 70.6 | **45.1** |
|  | 3 | 86,708 | 35,721,283 | 40.7 | 19.5 |
|  | 5 | 53,314 | 20,582,022 | 24.1 | 22.7 |
|  | 10 | 20,176 | 8,997,030 | 8.4 | 32.6 |
| **Scaffold28**   - **4.8Mbps,** - **# Gaps: 244** - **# N: 243.5Kbps** | 1 | 168,527 | 64,705,613 | 76.1 | 35.8 |
|  | 3 | 86,588 | 35,488,861 | 36.5 | 37.4 |
|  | 5 | 54,845 | 23,042,400 | 18.5 | **71.3** |
|  | 10 | 22,122 | 9,720,551 | 9.4 | 59.8 |
| **Scaffold36**   - **4.3Mbps,** - **# Gaps: 332** - **# N: 402.3Kbps** | 1 | 146,548 | 56,652,569 | 50.6 | **66.9** |
|  | 3 | 72,373 | 29,750,862 | 31.8 | 35.3 |
|  | 5 | 44,868 | 18,949,958 | 17.9 | 62.6 |
|  | 10 | 17,153 | 7,577,638 | 7.6 | 50.6 |
| **Scaffold39**   - **4.2Mbps,** - # Gaps: 387 - **# N: 505.1Kbps** | 1 | 134,141 | 52,281,320 | 35.4 | 25.2 |
|  | 3 | 62,129 | 25,699,474 | 28.9 | 20.7 |
|  | 5 | 36,790 | 15,609,672 | 13.4 | **56.6** |
|  | 10 | 13,024 | 5,735,802 | 4.7 | 38.5 |

**Supplementary Table 3. Scaffold-based gap filling analysis on scaffold 21, 28, 36 and 39 using non-duplicate read pairs**

|  | **# of unique pairs** | **#Recruited Barcode** | **#Recruited Read Pairs** | **Wall Clock Time (hr)** | **Filled Gap Length (%)** |
| --- | --- | --- | --- | --- | --- |
| **Scaffold21**   - **5.2Mbps,** - # Gaps: 313 - **# N: 383.3Kbps** | 1 | 177,302 | 64,370,474 | 74.1 | **51.8** |
|  | 3 | 86,413 | 33,744,148 | 31.5 | 47.8 |
|  | 5 | 53,087 | 20,893,778 | 22.3 | 46.1 |
|  | 10 | 20,018 | 8,259,223 | 7.0 | 38.0 |
| **Scaffold28**   - **4.8Mbps,** - **# Gaps: 244** - **# N: 243.5Kbps** | 1 | 167,131 | 60,358,361 | 62.3 | **79.9** |
|  | 3 | 86,336 | 32,925,945 | 31.6 | 70.6 |
|  | 5 | 54,597 | 21,313,213 | 20.9 | 65.5 |
|  | 10 | 21,965 | 8,929,363 | 9.6 | 56.6 |
| **Scaffold36**   - **4.3Mbps,** - **# Gaps: 332** - **# N: 402.3Kbps** | 1 | 145,305 | 52,877,734 | 48.1 | **65.9** |
|  | 3 | 72,176 | 27,639,712 | 26.7 | 59.4 |
|  | 5 | 44,677 | 17,541,650 | 16.5 | 58.5 |
|  | 10 | 17,018 | 6,955,085 | 8.4 | 49.8 |
| **Scaffold39**   - **4.2Mbps,** - # Gaps: 387 - **# N: 505.1Kbps** | 1 | 133,138 | 48,774,109 | 34.0 | 59.5 |
|  | 3 | 61,929 | 23,840,394 | 21.1 | **60.1** |
|  | 5 | 36,602 | 14,412,116 | 11.0 | 53.8 |
|  | 10 | 12,914 | 5,257,810 | 5.6 | 36.7 |

**Supplementary Table 4. Gap filling analysis for each partition algorithm**

| **Partition** | Scaffold-based | | | | **Gap-based** | | | | Window-based | | | |
| --- | --- | --- | --- | --- | --- | --- | --- | --- | --- | --- | --- | --- |
| **Scaffold ID** | Scaf21 | Scaf28 | Scaf36 | Scaf39 | Scaf21 | Scaf28 | Scaf36 | Scaf39 | Scaf21 | Scaf28 | Scaf36 | Scaf39 |
| **#Task** | 1 | 1 | 1 | 1 | 311 | 243 | 332 | 386 | 69 | 58 | 55 | 55 |
| **#Total Processed Barcode** | 86,707 | 86,588 | 72,373 | 62,128 | 82,837 | 66,143 | 89,343 | 86,289 | 70,280 | 62,097 | 63,542 | 54,707 |
| **#Total Processed Read Pair** | 35,721,245 | 35,488,861 | 29,750,862 | 25,699,346 | 35,181,566 | 27,996,758 | 37,579,213 | 36,519,860 | 29,524,954 | 25,985,365 | 26,489,125 | 22,908,558 |
| **Wall Clock Time**  **(hr)** | 45 | 37 | 32 | 18 | 5.9 | 6.2 | 4.1 | 6.3 | 2.3 | 2.0 | 2.1 | 2.0 |
| **Filled Gap Length (%)** | 19.5 | 37.4 | 35.3 | 20.7 | **65.0** | **80.2** | **76.8** | **68.5** | 57.4 | 68.8 | 71.6 | 60.4 |

**Supplementary Table 5. Analysis of filled gaps by LOCLA on LLD0021C**

|  |  | **Supernova** | **LOCLA** |
| --- | --- | --- | --- |
| Genome size |  | 2,894,333,848 | 3,143,552,207 |
| Total # of gaps |  | 64,987 | 55,210 |
| # of filled gaps |  | 0 | 9,777 |
|  | Completely-filled | 0 | 5,785 |
|  | Partially-filled | 0 | 3,992 |
| Filled gap count % |  | 0 | 15.04% |
| Total gap length (bp) |  | 274,647,410 | 256,926,360 |
| Filled gap length (bp) |  | 0 | 17,721,050 |
| Filled gap length % |  | 0 | 6.45% |
| **Average Filled Gap Length (bp)** |  | 0 | **1,812.52** |
|  | Completely-filled | 0 | **113.73** |
|  | Partially-filled | 0 | **4,274.27** |
| **Maximum filled gap length (bp)** | Completely-filled | 0 | **59,608** |
|  | Partially-filled | 0 | **100,000** |

**Supplementary Table 6. LOCLA outperforms Supernova in repeat sequences (sample: LLD0021C)**

| **Region** | | **Supernova Length** | | **LOCLA Length** | **Increased Percentage** |
| --- | --- | --- | --- | --- | --- |
| **SINEs:** | 360,307,214 | | 367,599,283 | | 2.02% |
| ALUs | 297,675,946 | | 304,339,130 | | 2.24% |
| MIRs | 62,062,069 | | 62,685,814 | | 1.01% |
| **LINEs:** | 555,422,256 | | 570,384,310 | | 2.69% |
| LINE1 | 467,877,986 | | 481,834,046 | | 2.98% |
| LINE2 | 78,456,751 | | 79,370,127 | | 1.16% |
| L3/CR1 | 7,146,074 | | 7,219,478 | | 1.03% |
| **LTR elements:** | 235,284,651 | | 241,576,774 | | 2.67% |
| ERVL | 48,026,801 | | 49,125,199 | | 2.29% |
| ERVL-MaLRs | 97,217,164 | | 99,198,073 | | 2.04% |
| ERV_classI | 78,440,079 | | 80,985,565 | | 3.25% |
| ERV_classII | 8,176,299 | | 8,798,235 | | 7.61% |
| **DNA elements:** | 86,856,844 | | 88,266,581 | | 1.62% |
| hAT-Charlie | 36,794,256 | | 37,280,601 | | 1.32% |
| TcMar-Tigger | 32,699,038 | | 33,364,831 | | 2.04% |
| **Unclassified:** | 4,164,825 | | 4,329,007 | | 3.94% |
| Small RNA | 1,171,956 | | 1,235,812 | | 5.45% |
| Satellites | 9,122,975 | | 14,774,390 | | 61.95% |
| Simple repeats | 36,143,467 | | 39,011,022 | | 7.93% |
| Low complexity | 6,216,364 | | 6,361,304 | | 2.33% |

**Supplementary Table 7. Analysis of filled gaps by LOCLA on CHM13**

|  |  | **Supernova** | **LOCLA** |
| --- | --- | --- | --- |
| Genome size (bp) |  | 2,914,622,463 | 2,951,434,376 |
| Total # of gaps |  | 25,814 | 7,178 |
| # of filled gaps |  | 0 | 18,636 |
|  | Completely-filled | 0 | 10,768 |
|  | Partially-filled | 0 | 7,868 |
| Filled gap count % |  | 0 | **72.19%** |
| Total gap length (bp) |  | 47,768,807 | 7,465,120 |
| Filled gap length (bp) |  | 0 | 40,303,687 |
| Filled gap length % |  | 0 | **84.37%** |
| **Average Filled Gap Length (bp)** |  |  | **2162.679062** |
|  | Completely-filled | 0 | **231.8** |
|  | Partially-filled | 0 | **4,805.00** |
| Maximum filled gap length (bp) | Completely-filled | 0 | **100,000** |
|  | Partially-filled | 0 | **100,000** |

**Supplementary Table 8. Gap closure results employing SOAPdenovo (GapCloser) and GapFiller obtained on Homo sapiens (chromosome 14)**

| **chromosome 14** | **Gap Filling Methods** | | |
| --- | --- | --- | --- |
|  | **Original assembly** | **SOAPdenovo (GapCloser)** | **GapFiller** |
| Genome size (bp) | 95,081,274 | 95,059,687 | 95,072,801 |
| Scaffolds | 19,249 | 19,249 | 19,249 |
| Gap count | 2,820 | 1,986 | 1,682 |
| Closed gap count % | 0 | 29.57% | 40.35% |
| Total gap length (bp) | 949,137 | 423,107 | 699,550 |
| Closed gap length % | 0 | 55.42% | 26.30% |
| **Average closed gap length (bp)** | **0** | **264.87** | **148.39** |

**Supplementary Table 9. LOCLA is capable of reducing a considerable amount of gaps through several iterations**

| **Stages of eel genome assembly** | **initial** | **GCB Gap Filling 1st iteration** | **GCB Gap Filling 2nd iteration** | **GCB Gap Filling 3rd iteration** |
| --- | --- | --- | --- | --- |
| Number of Scaffolds | 4,617 | 4,617 | 4,617 | 4,617 |
| Average Scaffold Length (bp) | 234,776 | 241,403 | 243,673 | 248,489 |
| Minimum Scaffold Length (bp) | 889 | 889 | 889 | 889 |
| Maximum Scaffold Length (bp) | 73,940,153 | 74,964,264 | 75,377,128 | 76,365,690 |
| N50 (bp) / L50 | 42,610,991 / 10 | 43,649,922 / 10 | 44,030,935 / 10 | 44,552,354 / 10 |
| N75 (bp) / L75 | 25,879,373 / 17 | 25,279,375 / 18 | 25,629,905 / 18 | 26,379,933 / 18 |
| N99 (bp) / L99 | 9,268 / 889 | 10,134 / 881 | 10,368 / 872 | 10,781 / 869 |
| Total bases in scaffolds (bp) | 1,083,962,265 | 1,114,555,594 | 1,125,040,273 | 1,147,271,934 |
| Total number of N | 47,921,917 | 19,671,117 | 15,111,710 | 12,579,893 |
| Reduced number of N | 0 | 28,250,800 | 4,559,407 | 2,531,817 |
| N % | 4% | 1.76% | 1.34% | 1.10% |
| Total bases without N (bp) | 1,036,040,348 | 1,094,884,477 | 1,109,928,563 | 1,134,692,041 |
| Increased bases without N (bp) | 0 | 58,844,129 | 73,888,215 | 98,651,693 |

**Supplementary Table 10. Hardware configuration and software information**

|  | **Server A** | **Server B** |
| --- | --- | --- |
| Architecture: | x86_64 | x86_64 |
| CPU op-mode(s): | 32-bit, 64-bit | 32-bit, 64-bit |
| CPU(s): | 96 | 160 |
| Memory: | 1511GB | 2267GB |
| Model name: | Intel(R) Xeon(R) CPU E7-4830 v3  @ 2.10GHz | Intel(R) Xeon(R) Gold 6230 CPU  @ 2.10GHz |
| Software: | Ubuntu 16.04.7 | Ubuntu 20.04 |

**Supplementary Table 11. The runtime of each LOCLA module on the 96-core CPU server (sample: LLD0021C)**

| **Module** | **Elapsed Time (d/ h:m:s)** |
| --- | --- |
| GCB Gap Filling | 1:41:37 |
| GCB Scaffolding | 18:30:11 |
| LCB Gap Filling | 22d/ 4:21:23 |
| LCB Scaffolding | 13d/17:02:57 |

**Supplementary Table 12. The runtime of LCB Gap Filling and LCB Scaffolding on the 96-core CPU server (sample: LLD0021C)**

| **Module** | **Process** | **Elapsed Time (d/ h:m:s)** |
| --- | --- | --- |
| LCB Gap Filling | Collect barcodes for each gap | 2:42:59 |
|  | Assemble contigs for each gap | 22d/ 3:6:18 |
|  | Fill in gaps | 1:32:06 |
| LCB Scaffolding | Assemble contigs for each scaffold pair | 11d/ 1:12:27 |
|  | Connect scaffolds | 2d/ 15:50:30 |

**Supplementary Table 13. Reducing runtime by distributing assembling tasks on 8 Virtual Machines (sample: LLD0021C)**

| **Strategy** | **Hardware Configuration** | **Total processed Read Pair** | **Elapsed Time**  **(d/ h:m)** | **Tasks per run** |
| --- | --- | --- | --- | --- |
| Parallel on 8  Azure VMs | 576 vCPU,  1,152 GB RAM | 2,141,896,476 | 15:27 | 14 |
| Sequentially on one server | 96 CPUs,  1511GB RAM | 3,254,750,449 | 22d/ 03:06 | 32 |

**Supplementary Table 14. Runtime of assembling contigs (LCB Gap Filling) for LLD0021C and CHM13**

| **Sample** | **Hardware Configuration** | **# of total processed Read Pairs** | **# of jobs in parallel** | **Elapsed Time (d/h:m:s)** |
| --- | --- | --- | --- | --- |
| LLD0021C | **96** CPUs, 1511GB RAM | 3,551,946,069 | **8** | **21**/06:19:42 |
| CHM13 | **160** CPUs, 2267 GB RAM | 3,535,752,766 | **16** | **5**/4:40:17 |

**Supplementary Table 15. LOCLA increased the content of disease-related genes on the Supernova assembly (sample: LLD00021C)**

| **Gene**  **Name** | **Function** | **Related Disease** | **Interaction Gene** | **Paralog** | **Total sequence**  **filled in gene (bp)** |
| --- | --- | --- | --- | --- | --- |
| **DDX11** | DNA helicase, DNA replication,  DNA repair, Genome stability, Spliceosome, | Bladder cancer,  Gastric Cancer, Melanoma, Warsaw syndrome, Fanconi anemia | SMC1A, STAG1, STAG2, CHTF18, DDX11-AS1 | BRIP1, RTEL1, RTEL1-TNFRSF6B, ERCC2 | 30,947 |
| **DDX11-AS1** | ncRNA, Increase DDX11  ATPase activity, Oncogenic driver | Glioma, Gastric cancer, Colorectal cancer, Brain tumor, Bladder cancer | DDX11 |  | 3,533 |
| **SMC1A** | Sister chromatid cohesion,  DNA repair, ATM/ATR regulation, G1/S checkpoint | Breast cancer, Colorectal cancer, Prostate cancer,  Cornelia de Lange syndrome, encephalopathy | SMC3, SMC1B, BRCA1, ATM, ATR | SMC1B | 84 |
| **STAG1** | Sister chromatid cohesion,  Spindle pole assembly, Meiosis, CDK-mediated phosphorylation | Colorectal cancer, Cornelia de Lange syndrome, Mental retardation | SMC1A, SMC1B, SMC3, RAD21, STAG2, STAG3 | STAG2 | 4,967 |
| **STAG2** | Sister chromatid cohesion,  Sister chromatid separation, CDK-mediated phosphorylation Aneuploidy | Breast cancer,  Sarcoma, Leukemia, Bladder cancer, Prostate cancer, MKMS syndrome | SMC1A, SMC1B, SMC3, RAD21, STAG1, STAG3 | STAG1, STAG3, STAG3L3, STAG3L4 | 770 |
| **CHTF18** | Sister chromatid cohesion,  Replication factor, Gastric cancer network  S phase checkpoint | Gastric cancer,  Cornelia de Lange syndrome | CTF18, CTF8, DCC1, RFC2, RFC3, RFC4, RFC5, PCNA, POLH |  | 143 |
| **RTEL1** | Double strand break repair,  Strand exchange, Holiday Junction formation, Telomere protect,  Anti-recombinase | Gastric cancer, Lung cancer,  Glioma, Pulmonary fibrosis, Dyskeratosis congenita | TERF1, PCNA, MMS19, GTF2H4, GTF2H1, MNAT1 | DDX11,  RTEL1-TNFRSF6B | 270 |

**Supplementary Table 16. Genes revealed after enhancing Supernova by the LOCLA assembly (sample: CHM13)**

| **Gene Name** | **Function** | **Related Disease** | **Related Pathway** | **Interaction Gene** | **Paralog** | **Total sequence**  **filled in gene (bp)** |
| --- | --- | --- | --- | --- | --- | --- |
| SPATA31C1 | Cell differentiation,  Spermatogenesis | Head and Neck Cancers | - | - | SPATA31C2, DKFZp434M131, SPATA31B1, SPATA31A2, SPATA31A2, FAM75A2, SPATA31A7, SPATA31A6, SPATA31A5, SPATA31A4, SPATA31A3 | 866 |
| DSPP | Dentinogenesis | Dentinogenesis  Imperfecta 1,  Dentinogenesis Imperfecta Shields Type Iii,  Dentin Dysplasia  Type Ii,  Dfna39/Dentinogenesis Imperfecta 1 Syndrome,  Hereditary  Opalescent Dentin | ECM proteoglycans,  Degradation of the extracellular matrix | TGFB1, BMP6, RUNX2, SP7, NOTCH2,  NOTCH1, NOTCH4, WNT10A, BMP15,  ITGB1, NOTCH3, BMP7, MYLIP, BMP2, BMP10, TWIST1 | - | 459 |
| MUC3A | Epithelial glycoprotein,  Ligand binding,  Intracellular signaling | Cap Polyposis, Lung Mucoepidermoid Carcinoma,  Colorectal Cancer,  Inflammatory Bowel Disease | Hyperphosphatemic tumoral calcinosis (HFTC),  Diseases of glycosylation,  Innate Immune System,  Metabolism of proteins,  O-linked glycosylation of mucins | GALNT8, B3GNT6, B3GNT2, GALNT9,  GCNT1, GALNT4, ST3GAL2, ST6GAL1,  GALNT6, GCNT3, ST3GAL4, C1GALT1, B3GNT5, GALNT1, B4GALT5, GCNT4, | MUC17, MUC13 | 405 |
| ATXN3 | Deubiquitination | Machado-Joseph Disease (MJD),  Spinocerebellar Ataxia  Type 3 | Proteolysis Putative ubiquitin pathway Singleton,  Regulation of degradation of deltaF508 CFTR in CF | NEDD8, XRCC6, EP300, FOXOO4,  CDKN1A, NCOR1, TEX11, HDAC3, PSMD4 | ATXN3L | 87 |
| KRTAP5-7 | Keratin-Associated Protein Coding | Childhood Germ Cell Cancer, Peroxisome Biogenesis Disorder 2a (PBD2A),  Cornelia De Lange Syndrome | Keratinization, development biology | KRTAP1-4, NME1-NME2,  KRT222, MICAL1, KT27, KRTAP4-6,  KRTAP3-2, KRTAP4-2, KRTAP4-7,  NCKIPSD, KRT32, KRT35, KRT39, KRT20,  KRT23, KRTAP1-1, KRT38, CASP14, KRTAP1-3, KRT31, KRT36, KRT34, KRT39 | - | 42 |
| ARHGAP11A-SCG5 | - | Hereditary Mixed Polyposis Syndrome | - | - | ARHGAP11A, ARHGAP11B | 18 |
| GDF7 | Prostate carcinoma,  Apolipoprotein B measurement,  Pelvic organ prolapse,  Uterine prolapse | Shoulder Impingement Syndrome,  Epicondylitis,  Endometrial Neoplasms | - | CHRD, BMP4, BMP2, SMAD2, BMP7,  NOG, SMAD1, BMP1, BMP15, BMPR2,  BMP10, SMAD5, NTF3, AMHR2 | GDF6, GDF5, BMP2, BMP4, BMP10, BMP7, BMP8B, NODAL, TGFB3, INHBE | 9 |
| ARHGAP11A | Cell-cycle arrest, Apoptosis | Chromosome 15q13.3 Deletion Syndrome,  Prader-Willi Syndrome 1,  Benign Epilepsy With Centrotemporal Spikes,  Colon cancers,  Human basal-like breast cancer cell line | p75 NTR receptor-mediated signaling, Signaling by GPCR | RHOD, RAC2, RHOA, CDC42, RHOB,  RHOC, RHOU, RHOQ, THAP1, RHOV | ARHGAP11A-SCG5, ARHGAP11B | 5 |
| OR2T35 | Odorant receptor | - | Signaling by GPCR | CUL3, GNAL, BAG3,  GNGT1, GNB1 | OR2T2, OR2T11,OR2T27, OR2T1, OR2T5 | 3 |
| PRB4 | Proline-Rich Salivary protein coding | Hypopharynx Cancer ,  Charcot Marie Tooth Disease | - | FURIN, S100A6 | PRB3, PRB2, PRB1, PRH1, PRH2 | 2 |
